# Direct Mapping of CDK2 Substrates in Embryonic Stem Cells Uncovers an AP-Site Repair Mechanism via HMCES Phosphorylation

**DOI:** 10.64898/2026.05.18.726044

**Authors:** Benjamin R. Topacio, Eli-Eelika Esvald, Jürgen Tuvikene, Lida Langroudi, Tapan K. Maity, Lisa M. Jenkins, Travis H. Stracker, Mardo Kõivomägi, S. Ali Shariati

**Author notes:** Corresponding authors ( &). Equal contribution (Benjamin R. Topacio, Eli-Eelika Esvald).

## Abstract

Embryonic stem cells (ESCs) proliferate rapidly while robustly maintaining genomic integrity and exhibiting high cell-cycle kinase activity. How this activity contributes to genome integrity remains unclear. Here, using mouse ESCs engineered to express an analog-sensitive CDK2, we combine thiophosphate labeling with mass spectrometry to define a high-confidence CDK2 substrate landscape. We uncovered 65 CDK2 substrates in total, including both known and previously unrecognized substrates. Among these, HMCES, a sensor of apurinic/apyrimidinic (AP) sites, was identified as a specific cyclin E-CDK2 substrate. We mapped three CDK2-dependent phosphorylation sites in HMCES and showed that phosphorylation of these sites decreased HMCES binding to ssDNA. Mutational analysis further revealed that HMCES docks to cyclin E-CDK2 complexes via the hydrophobic patch on cyclin E. Finally, we demonstrated that HMCES phosphorylation contributes to AP-site repair and promotes ESC proliferation. Together, our findings uncover a CDK2-HMCES signaling axis that links rapid cell-cycle progression to the preservation of genome stability in mouse ESCs.

## Main

Mouse embryonic stem cells (mESCs) can self-renew to unlimited numbers in culture while maintaining the potential to differentiate into any cell type in the adult body, a property known as pluripotency^1,2^. A defining feature of pluripotent mESCs is their restructured, accelerated division cycle, characterized by a shortened G1 phase and persistently high activity of the central cell cycle kinase, cyclin-dependent kinase 2 (CDK2)^3–7^. Several studies suggest that the high CDK2 activity of mESCs is linked to the pluripotency program and ability to maintain high fidelity DNA replication while undergoing rapid cell division^6,8–13^. However, a direct CDK2 substrate network and its functional relevance in mESCs have not been defined.

To address this, we engineered an analog-sensitive (AS) CDK2 in mESCs and determined the CDK2 substrate network by direct labeling. This approach identified both previously reported and novel CDK2 substrates. Strikingly, one of the most enriched functional groups in the CDK2 substrate network comprised proteins involved in DNA repair, supporting a direct role for CDK2 in maintaining genome stability in mESCs. Specifically, we identified HMCES as a direct substrate of CDK2. HMCES is a critical sensor of apurinic/apyrimidinic (AP) sites, among the most common DNA lesions that arise spontaneously or through DNA damage^14,15^. We mapped three phosphorylation sites on HMCES targeted by CDK2 and found that CDK2-mediated phosphorylation regulated binding of HMCES to DNA in vitro. Functional analysis showed that CDK2-mediated phosphorylation is essential for maintaining AP site repair capacity in the genome, and disruption of the CDK2-mediated phosphorylation of HMCES slowed mESC proliferation. Our findings uncover a previously unrecognized role of CDK2 in mediating a DNA repair quality-control system in rapidly dividing ESCs, providing insight into how CDK activity safeguards genome stability and early development.

## Results

### Optimization of analog-sensitive CDK2 system in mESCs

To study the function of CDK2 in mESCs (Fig. 1a) and enable direct identification of CDK2 substrates in mESCs, we incorporated the analog-sensitive mutation F80G, originally characterized in human CDK2^16–19^, into mouse CDK2. This mutant analog-sensitive CDK2 can specifically label substrates using bulky ATP-γ-S analogs. Thiophosphorylated substrates can then be labeled with p-Nitrobenzyl mesylate (PNBM) and analyzed by various methods using a thiophosphate ester-specific antibody (Fig. 1b)^20,21^.

**Figure 1.**
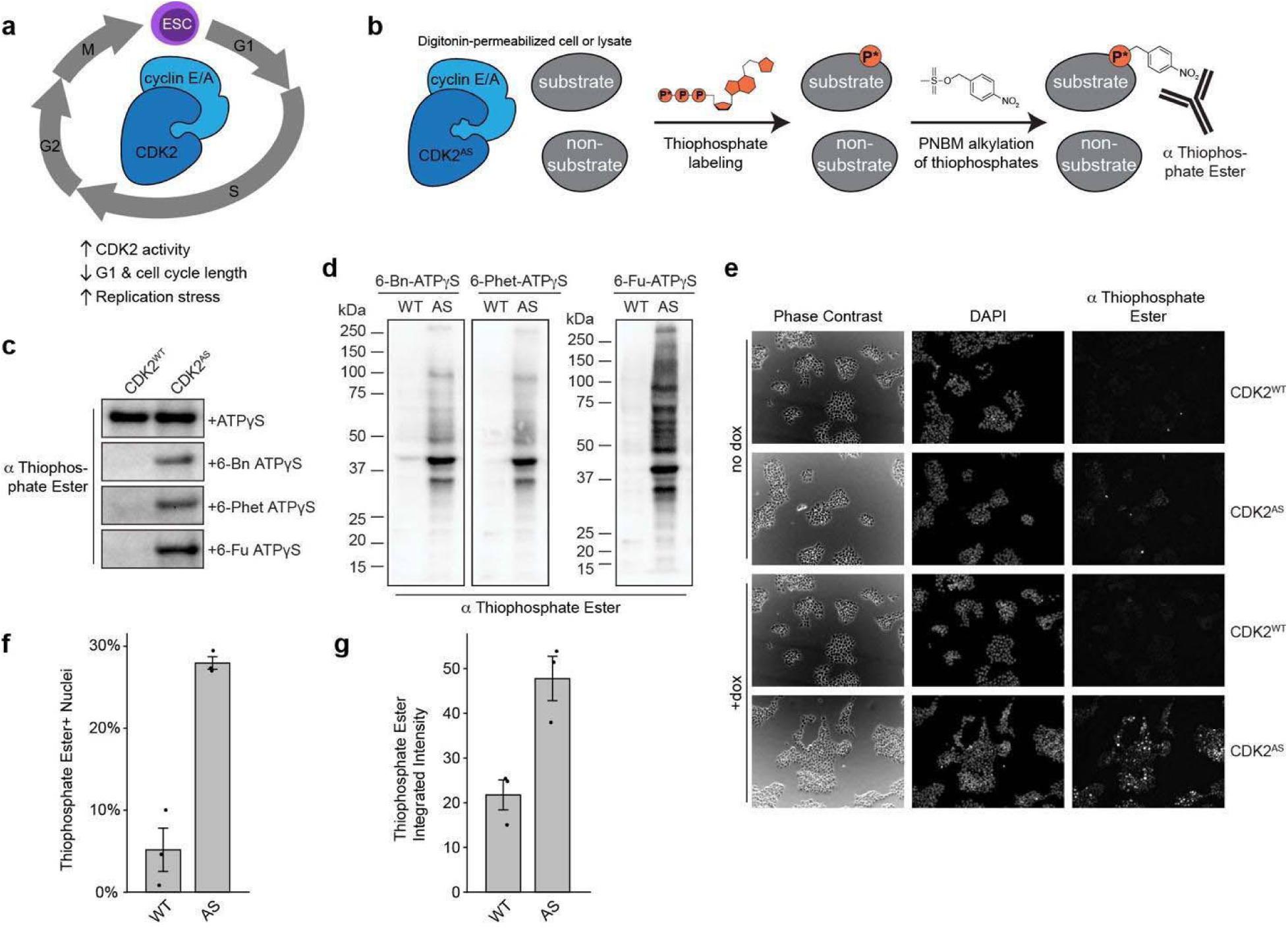
Establishment and functional validation of an analog-sensitive CDK2 system in mESCs. **a**, Schematic representation of the unique properties of the cell cycle in embryonic stem cells (ESCs). **b**, Schematic workflow of substrate labeling with analog-sensitive CDK2 (CDK2^AS^) and detection using a thiophosphate-specific antibody for immunofluorescence (IF), western blot (WB), or immunoprecipitation followed by mass-spectrometry (IP-MS). **c**, In vitro kinase assay with wild-type or analog-sensitive CDK2 (CDK2^WT^, CDK2^AS^) enriched from respective transgenic mouse ESCs and incubated with histone H1 model substrate and ATP-γ-S or the indicated bulky ATP analogs. **d**, In-cell labeling of CDK2 substrates using different bulky-ATP-γ-S analogs. **e**, Representative thiophosphate ester IF staining without CDK2^WT^ or CDK2^AS^ induction (no dox) or after 24h of induction with doxycycline (+dox). **f**,**g**, Quantification of thiophosphorylated protein-positive nuclei (**f**) or nuclear integrated intensity in positive nuclei (**g**) of +dox IF experiments from panel E (mean ± SD, n = 3).

We established cell lines expressing doxycycline-inducible V5-tagged wild-type or analog-sensitive CDK2 (CDK2^WT^ or CDK2^AS^) using a PiggyBac transposase system^22^. To confirm specificity of labeling by CDK2^AS^, we performed in vitro thiophosphorylation assays with CDK2^WT^ and CDK2^AS^ (Fig. 1c), using kinases enriched from the respective transgenic mESC lines. Kinases were incubated with the histone H1 (H1) model substrate^23^ in the presence of ATP-γ-S or one of three bulky ATP-γ-S analogs: 6-Bn-ATP-γ-S, 6-PhEt-ATP-γ-S, or 6-Fu-ATP-γ-S. To detect thiophosphorylated H1, we alkylated the thiophosphates with PNBM and performed immunoblotting for the thiophosphate ester bond^21^. While CDK2^AS^ and CDK2^WT^ had comparable activities using ATP-γ-S, only CDK2^AS^ was able to use the three bulky ATP-γ-S analogs to phosphorylate H1 (Fig. 1c), demonstrating specificity of the analog-sensitive mutation with this approach.

To optimize direct substrate labeling in cells, we determined the timing of CDK2^AS^ and CDK2^WT^ expression in response to doxycycline. We observed peak expression after eight hours of doxycycline induction, and high CDK2 levels were maintained for at least 24h without markedly affecting cell-cycle profiles (Extended Data Fig. 1a-b). We then compared thiophosphorylation in cells with different bulky ATP-γ-S analogs at different concentrations (Fig. 1d and Extended Data Fig. 1c). We observed high thiophosphorylation levels in CDK2^AS^ lines and only minor background levels in CDK2^WT^ lines. Specifically, we observed robust labeling with 100 µM of each bulky ATP-γ-S, with only a modest increase in labeling at the higher 500 µM concentration (Extended Data Fig. 1c). Finally, to confirm functionality of the CDK2^AS^ line, we studied in situ thiophosphorylation using immunofluorescence (IF) in asynchronously cycling mESCs after 24h of CDK2 induction by doxycycline (Fig. 1e). CDK2^AS^-expressing mESCs had appreciably more thiophosphate positive nuclei and higher levels of labeling compared to CDK2^WT^-expressing mESCs (Fig. 1f,g). Taken together, our data shows that CDK2^AS^ is functional in mESCs and can directly label CDK2 substrates in a specific-manner without altering cellular behavior, a conclusion consistent with previous analog-sensitive kinase studies^16–18,24^.

### Mass-spectrometry identifies the CDK2 substrate network in mESCs

We performed three independent labeling experiments in mESCs expressing CDK2^AS^ or CDK2^WT^, followed by affinity enrichment for thiophosphorylated proteins and identification by liquid chromatography tandem mass spectrometry (LC-MS/MS). Sixty-five statistically significant (p < 0.05) CDK2 substrates were identified by comparing CDK2^AS^ enriched thiophosphorylated proteins with CDK2^WT^ as a negative control (Fig. 2a and Supplementary Table 1). Several of these substrates have been previously identified as CDK2 targets, supporting the specificity and reliability of our CDK2^AS^-based labeling approach. These include well-established cell cycle regulators such as cell division cycle 20 (CDC20)^25,26^, protein regulator of cytokinesis 1 (PRC1)^27,28^, Bora, an activator of the aurora A kinase^29,30^, nucleophosmin 1 (NPM1)^31–33^, ankyrin repeat domain 17 (ANKRD17)^34^, DNA helicase B (HELB)^35,36^, and DNA ligase 1 (LIG1)^37^.

**Figure 2.**
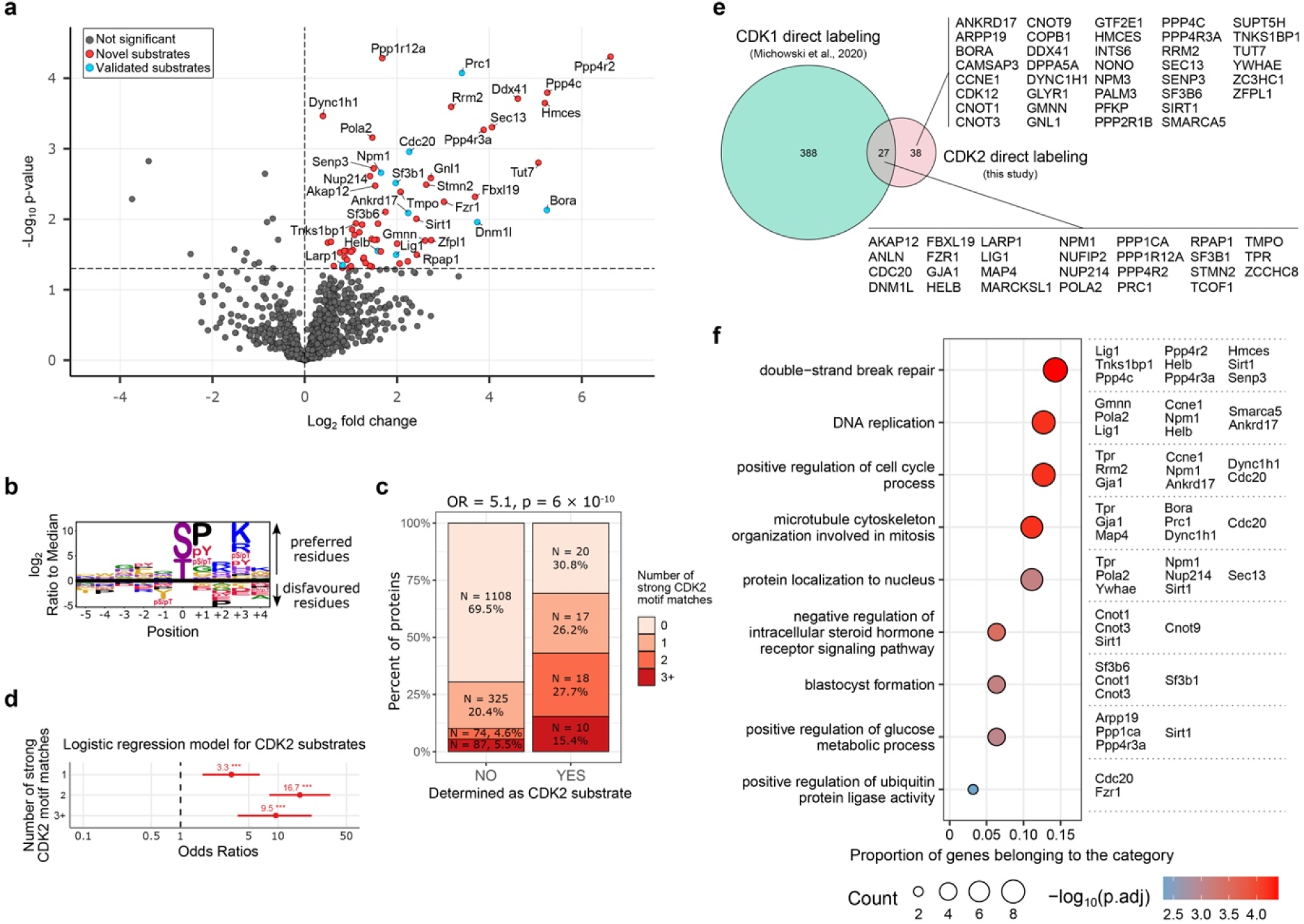
Identification of CDK2 substrates by chemical-genetic proteomics. **a**, Volcano plot showing enrichment of thiophosphorylated proteins in CDK2^AS^ vs CDK2^WT^-expressing cells as determined by LC-MS/MS. Proteins classified as CDK2 substrates were defined as those with log□ fold change (log□FC) > 0 and p < 0.05 in the mass-spec dataset (red dots). Blue dots represent the subset of identified CDK2 substrates that have been previously validated in the literature. Black dots indicate proteins not classified as CDK2 substrates. Full list of proteins is available in Supplementary Table 1. **b**, CDK2 phosphorylation site preference as described by Johnson et al., 2023^48^. **c**, Distribution of strong CDK2 consensus motif matches among proteins identified by mass spectrometry, grouped into CDK2 substrates and non-substrate controls. The odds ratio compares the odds of having at least one high-confidence CDK2 phosphorylation motif in CDK2 substrates versus non-substrate controls. **d**, Logistic regression model estimating the odds ratio of a protein being a CDK2 substrate as a function of the number of strong CDK2 motif matches. Proteins with zero strong CDK2 motif matches served as the reference group (odds ratio = 1). **e**, Venn diagram showing the overlap between CDK2 substrates identified in this study and CDK1 substrates previously identified in mESCs by direct labeling approach^24^. **f**, Overrepresentation analysis of functional categories among CDK2 substrates. The identified substrates belonging to the indicated category are shown on the right.

In addition to these canonical cell-cycle proteins, our analysis also identified several previously validated substrates involved in other cellular processes, including splicing factor 3b subunit 1 (SF3B1)^38–40^, La ribonucleoprotein 1 translational regulator (LARP1)^41^, and the mitochondrial fission regulator dynamin-related protein 1 (DRP1, encoded by *Dnm1l*)^42^. Notably, CNOT1 and CNOT3, components of mRNA deadenylase CCR4-NOT complex that are well-established as supporting stem cell pluripotency^43–46^, were also identified as CDK2 substrates. Thus, this in-cell CDK2^AS^ labeling strategy captures both established and previously unrecognized CDK2 substrates, expanding the known CDK2 signaling network and highlighting the broader functional scope of CDK2 activity beyond canonical cell-cycle regulation.

Next, we sought to determine whether the identified significantly enriched 65 CDK2 substrates are enriched for the preferred CDK2 phosphorylation motif compared with other proteins identified by mass spectrometry. For this analysis we used a position weight matrix generated by Johnson et al. (Fig. 2b), which largely recapitulated the canonical CDK2 phosphorylation motif^47–49^. In this motif, the phospho-acceptor serine or threonine residues are immediately followed by proline and, preferably, a charged amino acid such as lysine or arginine is at +3 position (S/TPxK/R). Proteins identified as CDK2 targets (YES column, Fig. 2c) were significantly more likely to contain at least one high-confidence CDK2 phosphorylation motif than proteins not significantly detected by mass spectrometry (NO column, Fig. 2c) (odds ratio (OR) = 5.1, p = 6 × 10□¹□), and this enrichment increased with the number of phosphorylation sites (Fig. 2d). Specifically, proteins with one high-confidence phosphorylation site showed modest enrichment for being identified as substrates (OR = 3.3), whereas those with multiple consensus sites were significantly more likely to be detected as substrates (OR = 16.7 for two sites and OR = 9.5 for three or more good context sites) (Fig. 2d). As there is no comprehensive analysis of CDK2 phosphorylation direct targets or sites in mESCs, we also compared our dataset with a published list of S/TP phospho-sites found in mESCs^50^. These analyses revealed that our CDK2 substrates (YES column, Extended Data Fig. 2a) are enriched for S/TP-phosphorylated proteins compared to proteins not determined to be a CDK2 target in our mass-spectrometric analysis [NO column; odds ratio (OR) = 2.6, p = 0.00031; Extended Data Fig. 2a]. Together, our chemical-genetic approach enabled the identification of 65 high-confidence CDK2 targets in mESCs which show strong enrichment for CDK2 phosphorylation motifs.

A chemical-genetic approach, emplying a workflow distinct from that in this study, was previously used to identify CDK2 substrates in HEK293 cells^17^. Comparing our identified targets with those reported in HEK293 cells, we found a minor overlap of 10 substrates (CNOT3, FBXL19, NPM1, NUFIP2, PPP1CA, PRC1, RRM2, SMARCA5, TCOF1, TPR), likely due to the different proteome composition and CDK2 regulation in the two cell types (Extended Data Fig. 2b). Direct analog-sensitive labeling has also been used to identify CDK1 substrates in mESCs^24^. We found substantial overlap between our CDK2 substrates and reported CDK1 substrates in mESCs, consistent with the fact that both kinases are activated by cyclin A, which also mediates substrate recruitment. In contrast, substrates uniquely identified for CDK2 may, at least in part, be recruited through cyclin E (Fig. 2e). Of these 38 unique CDK2 substrates, ANKRD17 and SF3B1 have been previously reported as cyclin E-specific substrates^34,40^. Finally, based on the functional enrichment analysis of the identified substrates, CDK2 is involved in several key biological processes, including DNA repair, replication, and chromosome segregation (Fig. 2f). These processes are consistent with previously published functions of CDK2 in stem cells^51^ through molecular mechanisms that are not fully defined. Importantly, identification of substrates enriched for DNA repair and DNA replication pathways strongly implicates CDK2 in supporting faithful DNA replication and genome integrity.

### HMCES is a novel CDK2-specific substrate

Among the highest-confidence substrates for CDK2 was HMCES [5-hydroxymethylcytosine (5hmC) binding, ESC-specific], a DNA repair protein enriched in ESCs (Fig. 2a and Supplementary Table 1). HMCES has recently been described as a sensor and guardian of AP sites at the replication fork, where it helps to prevent fork stalling, mutations, and double-strand breaks^15,52–56^. Interestingly, despite their rapid cell cycle that is largely driven by constitutively high CDK2 activity^57,58^, pluripotent stem cells exhibit relatively low levels of DNA damage and higher expression of many DNA repair proteins, as compared to somatic cells^59–62^. We hypothesized that CDK2 supports faithful DNA replication during the rapid proliferation of stem cells by regulating AP site repair through HMCES and therefore selected it for further validation as a novel CDK2 substrate.

To validate HMCES as a CDK2 substrate and determine whether it is specifically phosphorylated by CDK2 or all cell cycle CDKs, we first treated mESCs with a panel of CDK inhibitors (CDKi), including PF-06873600 (CDK2/4/6-specific), INX-315 (CDK2-specific), PF-07104091 (CDK2-specific), RO-3306 (CDK1-specific), and Palbociclib (CDK4/6-specific), using a range of concentrations that block RB phosphorylation and induce cell-cycle arrest in different cell lines^63,64^. We analyzed HMCES phosphorylation with Phos-tag SDS-PAGE (hereafter referred to as Phos-tag; Fig. 3a and Extended Data Fig. 3a), which distinguishes phosphorylated HMCES from its non-phosphorylated counterpart. Interestingly, we only detected phosphorylated HMCES in control (DMSO-treated, asynchronously dividing) mESCs (Extended Data Fig. 3b). However, short treatment with CDK2 inhibitors, but not CDK1, CDK4, or CDK6 inhibitors, reduced HMCES phosphorylation (Fig. 3a and Extended Data Fig. 3b). Thus, HMCES is highly phosphorylated by CDK2 in mESCs.

**Figure 3.**
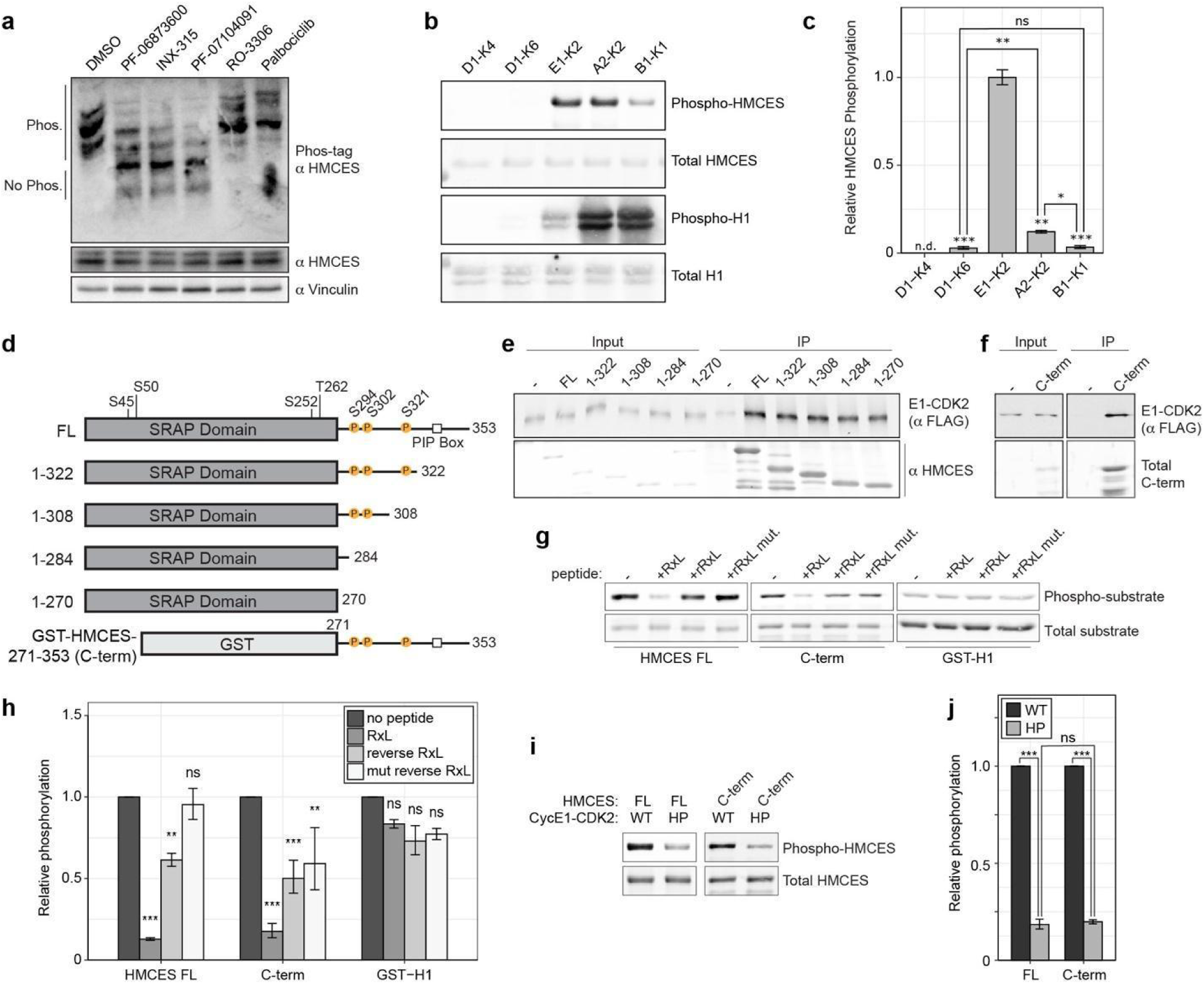
HMCES is a cyclin E-CDK2-specific substrate and its docking is predominantly mediated by the cyclin E’s hydrophobic patch. **a**, mESCs were treated with the indicated CDK inhibitors for 1h, lysed, and analyzed by Phos-tag or regular SDS-PAGE using anti-HMCES antibody or anti-Vinculin as loading control. **b**, In vitro kinase assay with different cell cycle cyclin-CDK complexes and full-length His-HMCES or histone 1 (H1) as a model substrate and incubated for 8 min at RT. Phosphorylation levels were assessed by Western blot using anti-thiophosphate ester antibody. Total protein levels were measured with Revert stain kit. **c**, Quantification of kinase assays shown on panel **b**. Data is shown relative to HMCES phosphorylation by cyclin E1-CDK2 (mean ± SEM, n = 3). **d**, Schematic representation of the generated truncated variants of HMCES. **e**,**f**, Co-immunoprecipitation analysis with purified His-HMCES fragments (**e**) or GST pull-down assay with C-terminal fragment of HMCES (C-term) (**f**) and FLAG-tagged cyclin E1-CDK2, analyzed by Western blot using anti-FLAG antibody. Equal amounts of HMCES variants were confirmed with anti-HMCES antibody or total protein stain in case of C-terminus of HMCES. **g**, In vitro kinase assay with full-length HMCES (HMCES FL), C-terminal fragment of HMCES (C-term), or GST-H1 and cyclin E1-CDK2 incubated for 5 min. The p21 peptide containing an RxL motif (denoted RxL), peptide containing a reverse RxL motif (rRxL), a peptide with mutated reverse RxL motif (rRxL mut), or buffer only as a control (denoted with a dash) were added to the reaction. **h**, Quantification of kinase assays shown on panel **g**. Data is shown relative to the control reaction with no added peptide for each substrate (mean ± SEM, n = 3). **i**, In vitro kinase assay with full-length HMCES (HMCES FL) or C-terminal fragment of HMCES (C-term) and wild-type or hydrophobic patch-mutated cyclin E1 linked to CDK2 (respectively WT or HP) incubated for 5 min. **j**, Quantification of kinase assays shown on panel **i**. Data is shown relative to each substrate phosphorylation level by wild-type cyclin E1-CDK2 (mean ± SEM, n = 3). n.s – p > 0.05, * – p < 0.05, ** – p < 0.01, *** – p < 0.001.

To confirm CDK-specificity of HMCES phosphorylation, we performed in vitro kinase assays using a panel of cyclin-CDK complexes representing distinct cell-cycle phases (Fig. 3b,c). Because CDK2 can be activated by either cyclin E or cyclin A, we tested both complexes. We observed a progressive increase in H1 phosphorylation from early G1 cyclin CDK complexes to complexes that are active later in the cell cycle (Fig. 3b), consistent with our previous findings^65^. In contrast, HMCES was not phosphorylated by cyclin D1-CDK4/6 complexes but was efficiently phosphorylated by cyclin E1-CDK2, cyclin A2-CDK2, and to a lesser extent by cyclin B1-CDK1 complexes (Fig. 3b). Comparison of HMCES phosphorylation with that of H1, a docking-independent substrate, revealed that cyclin E1-CDK2 exhibits the highest relative activity toward HMCES among the tested complexes, by approximately 10-fold (Fig. 3c). These findings confirm that HMCES is preferentially phosphorylated by CDK2 and suggests a specific molecular interaction between HMCES and cyclin E1-CDK2 complexes.

### The hydrophobic patch of cyclin E1 mediates the interaction with HMCES

To map the interaction sites between cyclin E1-CDK2 and HMCES, we first generated constructs that delete either the HMCES unstructured C-terminal domain or the conserved SOS response associated peptidase (SRAP) domain (Fig. 3d) and performed co-immunoprecipitation (co-IP) assays (Fig. 3e,f). Both the SRAP domain and the C-terminal fragment were able to bind cyclin E1-CDK2 complexes (Fig. 3e,f), consistent with a multivalent binding mechanism involving distinct docking regions, as has been reported for the interaction between SKP2 and cyclin A-CDK2 where two independent motifs on the SKP2 protein mediate binding to cyclin A^66^.

To determine the cyclin E interface that interacts with HMCES, we examined the cyclin E hydrophobic patch (HP) region. The HP region interacts with short linear motifs (SLiMs) that are often located within intrinsically disordered regions of substrates^67,68^. One of the best-characterized SLiMs that interact with HP regions on cyclins is the R/KxLF/LF/L motif (hereafter referred to as RxL)^69^. To test whether HMCES phosphorylation depends on HP-mediated interactions, we examined the effects of two competitor peptides on the interaction between cyclin E-CDK2 and HMCES. One peptide contains an RxL motif from the CDK inhibitor protein p21, which has been previously shown to bind cyclin E1^69^. Additionally, we examined a peptide containing a recently described reverse RxL motif, as well as a version with mutated interface residues as a control^66^. The p21-derived RxL peptide reduced phosphorylation of both full-length HMCES and the C-terminal fragment 6-8-fold, while the reverse RxL peptide decreased HMCES phosphorylation to a lesser extent (Fig. 3g,h). The reverse RxL peptide with mutated interface residues had no effect on full-length HMCES phosphorylation and only a minor effect on phosphorylation of the C-terminal fragment compared to control (∼1.7-fold decrease, Fig. 3g,h). Importantly, none of the peptides affected CDK2 kinase activity towards the docking-independent model substrate histone H1 (Fig. 3g,h). Finally, we tested whether the HP mediates cyclin E1-CDK2 docking of HMCES by directly mutating the HP region of cyclin E1 (Fig. 3i,j). As expected, mutations in the HP interface decreased phosphorylation of both full-length and the C-terminal fragment of HMCES approximately 5-fold, indicating that the cyclin E HP is the main docking site that contributes to the HMCES interaction with the cyclin E1-CDK2 complex. However, neither the RxL-containing peptide nor HP mutation on cyclin E fully abolished HMCES phosphorylation, suggesting that the HMCES-cyclin E1-CDK2 interaction might involve multiple docking regions, with the HP contributing significantly to this multivalent interaction.

### CDK2 phosphorylates the C-terminal half of HMCES

HMCES contains four potential CDK phosphorylation sites within the SRAP domain and three within the unstructured C-terminal region (Fig. 3d). To determine which of these sites are targeted by cyclin E1-CDK2, we measured the phosphorylation of truncated HMCES variants with quantitative in vitro kinase assays. Full-length HMCES, the C-terminal fragment, and all truncated variants were phosphorylated, except for the truncation lacking the C-terminus beyond amino acid 284 (named HMCES 1-284, Fig. 4a,b). To test the effect of these truncations in cells, we generated doxycycline-inducible mESC lines expressing either full-length or truncated HMCES variants and examined their phosphorylation level using Phos-tag. Consistent with the in vitro results, full-length HMCES was phosphorylated, whereas the largest truncation lacking residues beyond 284 was not (Fig. 4c). Thus, both in vitro and cellular assays indicate that the primary HMCES phosphorylation sites are located between amino acids 284-308.

**Figure 4.**
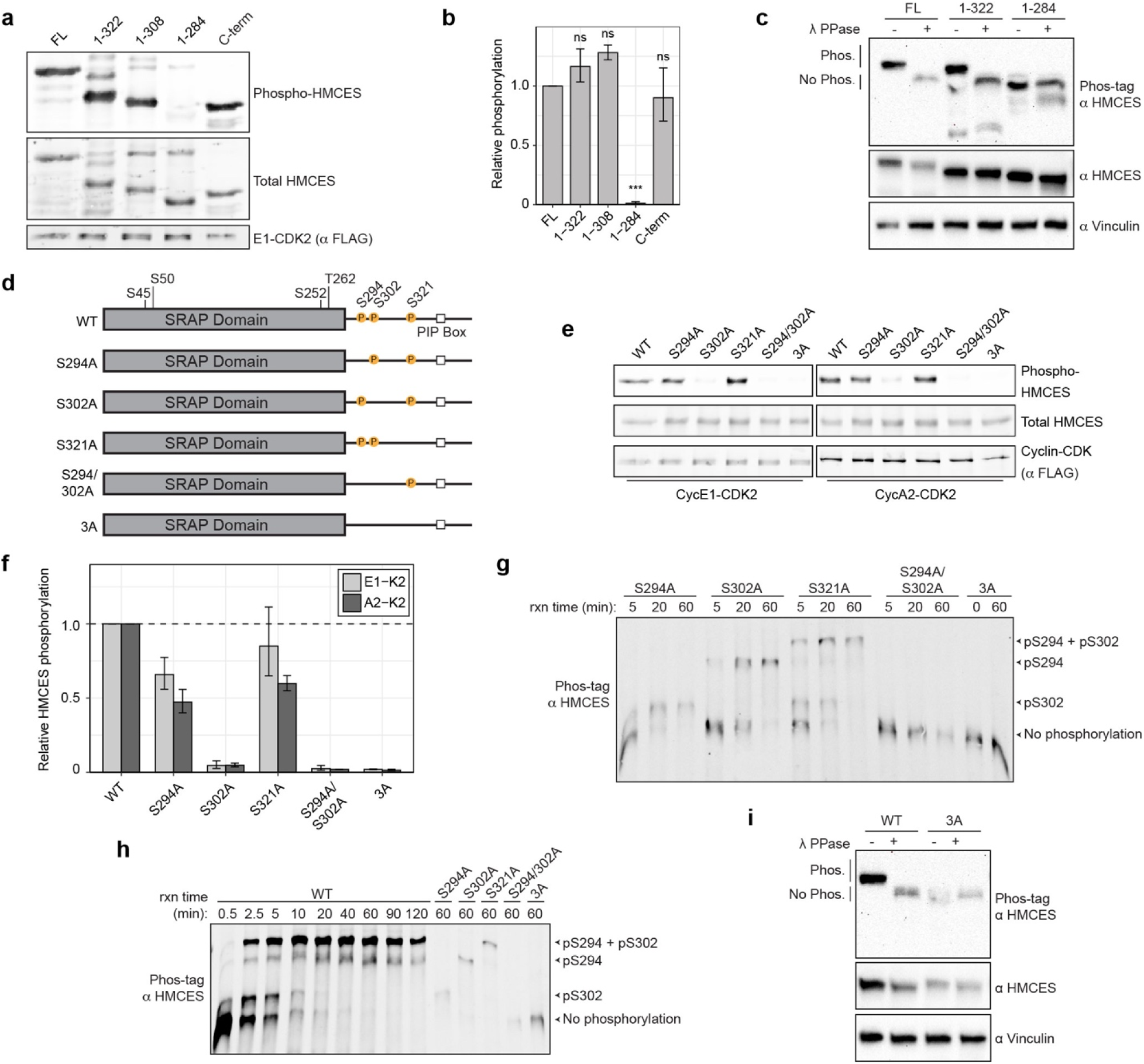
Cyclin E1-CDK2 phosphorylates sites within the C-terminal region of HMCES in a sequential manner. **a**, In vitro kinase assay with His-HMCES fragments or C-terminal fragment of HMCES (C-term) and cyclin E1-CDK2 incubated for 5 min. Phosphorylation levels were assessed by Western blot using anti-thiophosphate ester antibody and equal cyclin E1-CDK2 levels were confirmed with anti-FLAG antibody. Total protein levels were measured with the Revert stain kit. **b**, Quantification of kinase assays shown on panel **a**. Data is shown relative to full-length HMCES phosphorylation by cyclin E1-CDK2 (mean ± SEM, n = 4; n = 3 for 1-308 HMCES variant and n = 2 for GST-HMCES). **c**, Transgenic mESC cell lines expressing inducible Clover-tagged full-length HMCES or truncated variants were lysed and the lysates treated with λ phosphatase where indicated (λ PPase denoted with plus sign respectively) and analyzed by Phos-tag or regular SDS-PAGE using anti-HMCES or anti-Vinculin antibody as loading control. **d**, Schematic representation of the different serine to alanine phosphorylation-deficient HMCES variants used in **e**-**h**. **e**, In vitro kinase assay with full-length His-HMCES or HMCES phosphorylation variants incubated with cyclin E1-CDK2 or cyclin A2-CDK2 for 5 min. **f**, Quantification of kinase assays shown on panel **e**. Data is shown relative to wild-type HMCES phosphorylation by each cyclin-CDK complex (mean ± SEM, n = 2). **g**, In vitro kinase assay with mutated HMCES variants and cyclin E1-CDK2 for the time indicated were separated using Phos-tag. anti-HMCES antibody was used to follow HMCES phosphorylation. **h**, An in vitro time-course kinase assay with wild-type HMCES (HMCES WT) and cyclin E1-CDK2 over the indicated time points followed by separation using Phos-tag gels. HMCES phosphorylation was monitored by Western blot using anti-HMCES antibody, with phosphosite mutants included for comparison. **i**, Transgenic mESC cell lines expressing inducible Clover-tagged wild-type or triple alanine mutant HMCES (3A) and analyzed as in panel C. n.s – p > 0.05, * – p < 0.05, ** – p < 0.01, *** – p < 0.001.

We next sought to determine the contribution of each individual phosphorylation site between amino acids 284-308 to overall HMCES phosphorylation. The C-terminal region of HMCES contains three potential CDK phosphorylation sites at positions 294, 302, and 321. Of these sites, S294 and S302 have the canonical CDK consensus sequence (SPxK), whereas S321 resides within a suboptimal motif (SP). To determine the contribution of each site, we performed in vitro kinase assays with cyclin E1-CDK2, cyclin A2-CDK2, and cyclin B1-CDK1 complexes and several phosphosite variants: three single serine to alanine mutants, a double mutant targeting the two canonical sites (S294A/S302A), and a triple mutant in which all three sites were substituted with alanine (hereafter referred to as the 3A variant, Fig. 4d-f and Extended Data Fig. 4a,b). Mutation of all three sites (3A mutant) abolished phosphorylation by all tested cyclin-CDK complexes, while mutation of the suboptimal site (S321A) had a minor effect. In contrast, mutation of S302 markedly reduced phosphorylation (20-30-fold) by CDK2-containing complexes, whereas mutation of S294 had a moderate effect (∼2-fold; Fig. 4e,f). Altogether, these results indicate that HMCES is predominantly phosphorylated at S302, with additional contribution from S294, and to a lesser extent, S321.

The unexpected finding that only one of the two optimal CDK sites made a major contribution to overall phosphorylation suggests a coordinated site-specific phosphorylation mechanism. To examine the dynamics of individual phosphorylation events, we set up a HMCES phosphorylation time-course and analyzed samples using Phos-tag, which enabled us to resolve distinct phospho-species and monitor their kinetics. We first analyzed alanine-substituted HMCES variants to assign specific phosphosites to individual bands on Phos-tag. This approach confirmed that S294 and S302 represent the major phosphorylation signal (Fig. 4g). Time-course analysis revealed that phosphorylation at S302, as well as the double phosphorylated form (at S294 and S302) accumulated rapidly and reached a plateau within 40 min (Figure 4h and Extended Data Fig. 4c). In contrast, the single phosphorylated S294 species persisted after 2h of incubation with cyclin E1-CDK2. Finally, inducible expression of either wild-type or triple alanine-mutated HMCES in mESCs confirmed that these three sites account for all the HMCES phosphorylation in cells (Fig. 4i). Together, these findings suggest a phosphorylation model for HMCES with S302 functioning as a critical early phosphorylation site followed by phosphorylation of S294.

### CDK2-dependent HMCES phosphorylation regulates DNA repair and cell proliferation

Next, we asked whether CDK2-mediated phosphorylation regulates HMCES function in DNA repair and cell cycle progression. We first ruled out the effect of phosphorylation on HMCES abundance, as treatment with CDK inhibitors altered HMCES phosphorylation without affecting total HMCES protein levels (Fig. 3a). We therefore hypothesized that HMCES phosphorylation regulates its function.

The SRAP domain of HMCES is a well-established single-stranded DNA (ssDNA)-binding domain that mediates covalent attachment to DNA^15,56,70^. However, the phosphosites we identified are located in the disordered C-terminal region, outside of the SRAP domain^15,70^. This led us to first test whether the binding affinity of the SRAP domain differs from that of full-length HMCES using an electrophoretic mobility shift assay (EMSA, Fig. 5a,b). Strikingly, this revealed that the SRAP domain alone had a ∼10-fold lower affinity for ssDNA-binding affinity than full-length HMCES, suggesting that the C-terminal region is involved in its DNA binding-activity. To determine if phosphorylation of this region affects ssDNA binding, we incubated wild-type or phosphorylation-deficient HMCES 3A with cyclin E1-CDK2 in the absence (kinase active) or presence of EDTA (kinase inactive) for 1h and then performed EMSA. Phosphorylation decreased the affinity of wild-type HMCES for ssDNA ∼3-fold (Fig. 5c,d), whereas the 3A mutant was unaffected by incubation with cyclin E1-CDK2 (Fig. 5c,e), indicating CDK2-mediated phosphorylation reduces the affinity of HMCES for ssDNA. In addition, fluorescence polarization^70^ assays independently showed that HMCES binding to DNA was specifically impaired by phosphorylation (Fig. 5f). Taken together, our in vitro DNA binding assays demonstrate that phosphorylation of HMCES regulates its function by reducing its affinity for ssDNA.

**Figure 5.**
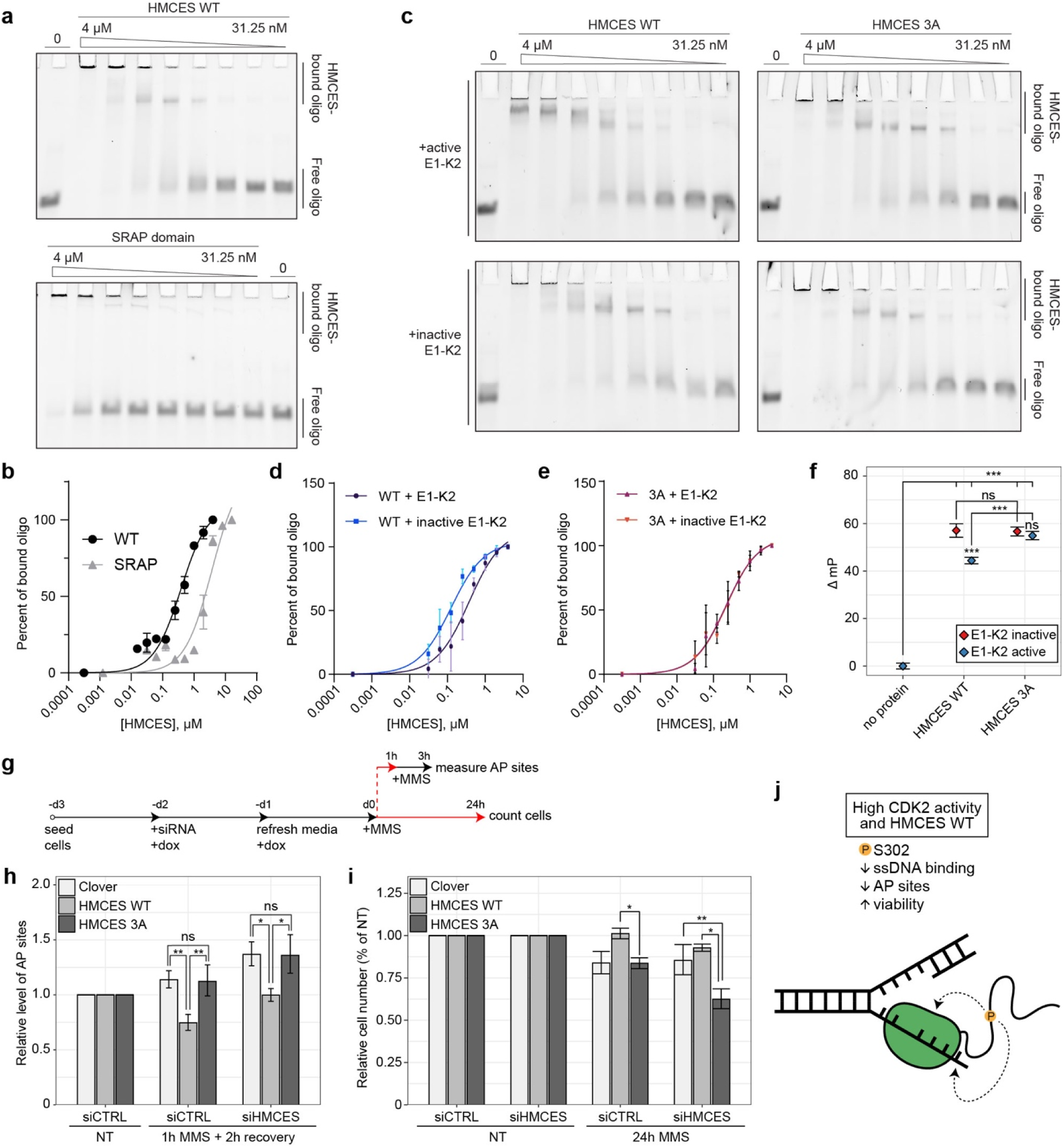
HMCES phosphorylation regulates its DNA binding, AP site repair, and cell viability. **a**, Electrophoretic mobility shift assay (EMSA) with full-length HMCES (HMCES WT) or SRAP domain of HMCES using 6FAM-labeled ssDNA probe. **b**, Quantification of panel **a** (n = 2). **c**, Representative image of EMSA with wild-type HMCES or triple alanine-mutant (3A) variant that were beforehand incubated with cyclin E1-CDK2 for 1h (denoted as E1-K2 active) or phosphorylation was prevented by adding EDTA to the reaction before the enzyme (denoted as E1-K2 inactive). **d**, Quantification of panel **c** HMCES WT data (n = 3). **e**, Quantification of panel **c** HMCES 3A data (n = 3). **f**, Fluorescence polarization assay using 6FAM ssDNA and HMCES WT and 3A variants incubated with cyclin E1-CDK2 as in panel **c**. Data is shown relative to the light polarization in samples without HMCES protein (mean ± SEM, n = 4). **g**, Schematic of AP site assay and cell viability assay experiments. Treatments to deplete endogenous HMCES with siRNA and express doxycycline-inducible siRNA-resistant wild-type HMCES (HMCES WT) or triple alanine-mutated HMCES (HMCES 3A) in transgenic mESC cell lines are the same for both AP site and cell viability assays. For the AP site assay, cells were treated with 10 mM MMS for 1hr, then MMS was removed, and cells were harvested after a 2h recovery period. For the cell proliferation assay, cells were grown in the presence of 2 mM MMS for 24h and then the number of viable cells were counted. **h**, Relative levels of AP sites in conditions described in **g**. Non-targeting siRNA (siCTRL) was used as a control for HMCES silencing, Clover induction served as a control for HMCES expression, and an equal volume of water was used as a no-treatment control (NT). Data are shown relative to AP-site level in the corresponding non-treated cells transfected with siCTRL (mean ± SEM, n = 5). **i**, Relative cell numbers after 24h of MMS. Data is shown relative to cell numbers in nontreated cells in respectively silenced and HMCES-expressing cells. (mean ± SEM, n = 5) n.s – p > 0.05, * – p < 0.05, ** – p < 0.01, *** – p < 0.001. **j**, The role of HMCES phosphorylation in mESCs. When CDK2 activity is high, HMCES is phosphorylated, the number of AP sites is lower, and cells divide rapidly. The C-terminal region is important for ssDNA binding and its phosphorylation modulates DNA binding by direct and/or indirect mechanisms.

HMCES knockout cells accumulate AP sites^15^. We therefore hypothesized that if HMCES binding is regulated by CDK2-mediated phosphorylation, then the overall burden of AP sites in mESCs might also be affected. To test this hypothesis, we expressed siRNA-resistant wild-type or 3A-mutant HMCES in mESCs where endogenous HMCES was siRNA-depleted (Fig. 5g) and induced AP sites with methyl methanesulfonate (MMS), as previously described^15^. Following depletion, siRNA-resistant HMCES wild-type and 3A were expressed at comparable levels (Extended Data Fig. 5a,b). AP site quantification with an aldehyde reactive probe^71^ showed that expression of wild-type HMCES reduced the number of AP sites, whereas the 3A mutant did not (Fig. 5g,h). MMS treatment also reduced cell proliferation, and this reduction was suppressed by expression of wild-type HMCES, but not 3A mutant (Fig. 5i). This effect is comparable to what has been observed in HMCES KO^55^ and HMCES mutant^53^ somatic cells in response to DNA damage. A similar effect was seen with potassium bromate treatment, another inducer of AP sites, although to a lesser extent (Extended Data Fig. 5c,d). Together, these cell-based assays demonstrate that CDK2-mediated phosphorylation of HMCES is critical for AP-site repair, likely by modulating the dynamics of HMCES-DNA binding, and that disrupting this phosphorylation impairs stem cell viability and proliferation (Fig. 5j).

## Discussion

Coordinated CDK2 activity is central for orderly progression of the cell cycle across different cell types^72^. In pluripotent ESCs, rapid cell division cycler driven by high CDK2 activity has also been linked to self-renewal, pluripotency programs and early lineage commitment decisions^4,51,57,73–76^. However, the downstream substrates through which CDK2 executes these functions in ESCs have remained poorly defined. Through a direct labeling approach, our work identified a high-confidence CDK2 substrate landscape in mESCs, composed of 65 candidate substrates of which only 18, to our knowledge, had been previously reported as CDK2 substrate. Furthermore, over 40% of the identified CDK2 substrates contain multiple consensus CDK2 phosphorylation sites, suggesting a broad capacity for multisite and hierarchical phosphorylation.

Specialized regulatory mechanisms in mESCs likely reflect unique requirements of the pluripotent state, where high constitutive CDK2 activity together with an ESC-specific proteome maintain rapid cell cycle progression and pluripotency. For example, CDK2 has been reported to phosphorylate core pluripotency factors OCT4, NANOG, and SOX2 in vitro^75^. Interestingly, these transcription factors were not detected in our mass-spectrometry experiments, possibly reflecting differences in assay sensitivities, growth conditions, or total levels of these proteins. Instead, among the pluripotency-related substrates, we identified the CCR4-NOT components, CNOT1, CNOT3, and CNOT9. CNOT1-3 are essential for maintaining mouse and human ESC identity and preventing extraembryonic differentiation^45^. Because the CCR4-NOT complex regulates mRNA translation and quality control^77^, our findings raise the intriguing possibility that CDK2 contributes to the pluripotency program through non-canonical pathways independently of its canonical cell cycle functions.

A key finding of our study is the identification of HMCES, a key sensor and stabilizer of AP sites^15,54^, as a direct and specific substrate of CDK2 in mouse ESCs. Our combined quantitative in vitro and in cell assays further indicate that HMCES is phosphorylated preferentially by cyclin E1-CDK2, whereas other cyclin-CDK complexes phosphorylate HMCES only weakly or not at all. This selectivity argues for a dedicated docking mechanism specific to cyclin E1, rather than CDK2-mediated recognition of a minimal phosphoacceptor motif. Indeed, our results implicate the hydrophobic patch region of cyclin E1 in recruiting HMCES predominantly through an interaction within its C-terminus. Interestingly, the C-terminus of HMCES does not contain any canonical [R/K]xL[F/L][F/L] sequences, suggesting that the cyclin E1 hydrophobic patch may, similarly to yeast cell cycle cyclins^69,78–80^, recognize alternative SLiMs. Because the existing structural studies have focused primarily on the isolated SRAP domain rather than full-length HMCES^70^, determination of a full-length HMCES-cyclin E1-CDK2 complex structure will be important for defining the mechanism of cyclin docking and selectivity.

Of the seven potential CDK phosphorylation sites in HMCES, our analysis indicates that CDK2 primarily targets S302, although all three phosphosites within the unstructured C-terminal tail contribute to overall HMCES phosphorylation. Compared with bacterial SRAP proteins, the C-terminal tail of eukaryotic HMCES appears to have expanded during evolution, enabling additional regulatory functions, including docking to PCNA through the PIP box^15^. Here, we report that the isolated SRAP domain has a substantially lower binding affinity for ssDNA compared to full-length HMCES, indicating that the C-terminal tail contributes to efficient DNA engagement. We further show that phosphorylation within the C-terminal tail reduces HMCES affinity for ssDNA. While it may seem paradoxical that CDK2-mediated phosphorylation weakens HMCES ssDNA binding, yet promotes AP-site repair, we propose that phosphorylation increases its dynamic behavior, preventing HMCES from becoming non-productively trapped on the abundant ssDNA present in stem cells^59^, allowing HMCES to more efficiently scan nascent DNA at replication forks for AP sites. In addition, CDK2 modulation of DNA binding could serve to prevent error prone microhomology-mediated end-joining (MMEJ) during S-phase^81–84^.

Because the DNA-binding activity of HMCES resides in the SRAP domain, whereas all three identified CDK2 phosphosites are located within the disordered C-terminus, several mechanisms could explain how phosphorylation modulates DNA binding. First, phosphorylation may increase electrostatic repulsion between HMCES and negatively charged DNA, thereby weakening ssDNA binding. Second, the negatively charged phosphorylated tail might result in conformational changes to partially block the HMCES DNA-binding surface, acting in an autoinhibitory manner. In addition, in cells, phosphorylation may alter interactions with proteins that bind the HMCES C-terminus which could indirectly alter DNA binding. Further studies are required to determine how CDK2-mediated phosphorylation of HMCES affects its structure and interactions with DNA and other proteins at the replication fork.

Despite HMCES’s evolutionary conservation from bacteria to mammals, the function and regulation of HMCES have remained understudied, and most mechanistic studies of its role in AP-site repair have been performed in cancer-derived or immortalized cell lines. In contrast, embryonic cells experience intrinsic replication stress due to their short G1 phase and persistently high CDK2 activity, representing a distinct physiological state to study HMCES regulation. Our findings suggest that in ESCs, CDK2-mediated phosphorylation of HMCES fine-tunes its DNA binding to ensure timely AP site repair and supporting faithful replication during rapid proliferative phase in early development, in agreement with higher HMCES expression levels in early embryos and HMCES knockout embryos exhibiting sublethal phenotype^85,86^.

## Methods

### Doxycycline-inducible PiggyBac integration vector construction

Mouse *Cdk2*-WT and *Cdk2*-F80G gBlocks were synthesized by Integrated DNA Technologies (IDT) and used as templates for PCR. Mouse *Cdk2* cDNA was then cloned into a PiggyBac integration plasmid with an N-terminal V5 tag, driven by the TRE3G doxycycline-inducible promoter. siRNA-resistant mouse *Hmces* cDNA was cloned into PiggyBac integration plasmids but with either no epitope tag or with a C-terminal 3xFLAG-Clover tag.

To generate siRNA-resistant mouse *Hmces* cDNA, silent mutations were introduced into sequences specifically targeted by the siGENOME Mouse Hmces (232210) siRNA SMARTpool (Horizon Discovery, #M-062972-00-0005) and gBlocks were synthesized by IDT. Reverse primers for *Hmces* PCR were designed to amplify full length or generate truncated versions of *Hmces*. To generate the 3A phosphorylation mutant *Hmces*, a full-length siRNA-resistant *Hmces* PiggyBac plasmid was digested with BspEI and EcoRI to remove a portion of the cDNA sequence comprising the C-terminal phosphosites. A gBlock representing the same fragment of *Hmces* with 3A substitutions was PCR-amplified, digested, and ligated into the digested plasmid to generate the siRNA-resistant *Hmces* 3A PiggyBac plasmid.

### Cell Culture

Mouse v6.5 embryonic stem cells (mESCs) were cultured in DMEM High Glucose, GlutaMAX (Gibco, #10566016) supplemented with 15% FBS (MilliporeSigma, #ES-009-B), 1000 U/ml LIF (MilliporeSigma, #ESG1107), 0.1 mM NEAA (Gibco, #11140050), 1 mM sodium pyruvate (Gibco, #11360070), 50 U/mL Penicillin-Streptomycin (Gibco, #15070063), and 0.1 mM 2-Mercaptoethanol (Gibco, #21985023). mESCs were grown on tissue culture-treated plates coated with 0.1% gelatin (Sigma G1890) in humidified incubators at 37°C, 5% CO_2_, and ambient O_2_.

Doxycycline-inducible *Cdk2* or *Hmces* variants were integrated into the genomes of mESCs by transfecting cells on a 6-well dish with 0.8 μg of doxycycline-inducible PiggyBac integration plasmid and 0.4 μg of PiggyBac Transposase plasmid (Ding et al., 2005) using the TurboFect Transfection Reagent (ThermoFisher R0531). Zeocin (final concentration 50 μg/mL; Gibco, #R25001) selection began two days after transfection with daily media changes and passaging as needed. Zeocin resistant cells were maintained as polyclonal cell lines and protein expression was induced with doxycycline (500 ng/mL) for one day or as specified in the experiment.

All CDK inhibitor experiments were performed at 37°C for 1h. Unless specified, the used concentrations were: 50 μM Palbociclib (PD-0332991; Selleckchem #S1116), 10 μM INX-315 (Selleckchem, #E1854), 50 μM RO-3306 (Cayman Chemical, #15149-5), 500 nM PF-06873600 (Cayman Chemical, #35502-5), and 50 μM PF-07104091 (Cayman Chemical, #41356).

### Immunoblotting

Cells were lysed in RIPA buffer supplemented with protease and phosphatase inhibitors. Protein concentration was determined using BCA Protein Assay kit (Pierce, #23227). Proteins from lysates were separated by SDS-PAGE on 10% polyacrylamide gels or precast 4-20% polyacrylamide gradient gels (Biorad, #4561095). To analyze phosphorylation, 10% polyacrylamide gels were supplemented with 100 μM MnCl_2_ (Sigma, #328146) and 50 μM Phos-tag Acrylamide (Fujifilm Biosciences, #AAL-107).

Proteins were transferred from gels to nitrocellulose membranes using the iBlot 2 dry blotting system (Invitrogen, #IB21001). Membranes were next incubated overnight at 4°C with the following antibodies: 1:2000 rabbit anti-Thiophosphate ester antibody [51-8] (Abcam, #ab92570), 1:1000 mouse anti-vinculin antibody (7F9): (Santa Cruz Biotechnologies, #sc-73614), 1:2000 rabbit anti-HMCES antibody (Atlas Antibodies, #HPA044968), 1:1000 mouse anti-FLAG M2 antibody (Sigma, #F1804), 1:1000 mouse anti-V5-Tag antibody SV5-Pk1 (Biorad, #MCA1360), and 1:2000 mouse anti-beta-tubulin (Sigma T5168). The primary antibodies were detected using goat anti-rabbit IgG H&L (HRP) (Abcam, #ab6721) and goat anti-mouse HRP (Azure Biosystems, #AC2115). Membranes were imaged on a Biorad and analyzed with Biorad Image Lab version 4.0.1

For phosphatase treated samples, cells were lysed in RIPA buffer supplemented with protease inhibitors only. After protein quantification, samples were combined with 1x PMP Buffer and 1 mM MnCl_2_ and incubated with Lambda protein phosphatase for 30 minutes at 30°C based on manufacturer’s specifications (NEB, #P0753S).

### Immunoprecipitation-coupled in vitro thiophosphorylation assays

V5-tagged CDK2 was purified from cell pellets using a V5-Trap® Agarose kit (ChromoTek, #v5tak) and eluted with 2 mg/ml V5 peptide (Sigma, #V7754) in 1X TBS. Equal amounts of purified enzyme were used for in vitro kinase assays at 30°C for 60 min with 2 μg of histone H1 (NEB, #M2501) in 50 mM HEPES pH 7.5, 150 mM NaCl, 5 mM MgCl_2_, 0.5 mM DTT, 0.2 mg/ml BSA, and 1 mM ATPγS (Abcam, #ab138911) or bulky ATPγS (6-Bn ATP-γ-S, Enzo, #BLG-B072; 6-PhEt ATP-γ-S, Enzo, #BLG-P026; 6-Fu ATP-γ-S, Enzo, #BLG-F008). Reactions were stopped by adding 20 mM EDTA (pH 8) and thiophosphate groups were alkylated by adding 2.5 mM PNBM (Selleckchem, #E1248) for 2h at RT with shaking. To stop alkylation, Laemmli buffer was added to final 1X, and samples were boiled for 5 min. Phosphorylated proteins were separated on 10% SDS-PAGE gels and phosphorylation was visualized using immunoblotting.

### In lysate thiophosphorylation

mESCs expressing CDK2 variants were lysed in 100 μL of lysis buffer (50 mM HEPES pH 7.5, 150 mM NaCl, 10 mM MgCl_2_, 0.4% NP-40) with protease inhibitors and cleared by centrifugation. Cleared lysates were supplemented with 2 mM TCEP, 3 mM GTP, 200 μM ATP, 40 nM creatine phosphate (CRPHO-RO) and 0.2 mg/ml creatine kinase (CK-RO) for ATP regeneration^87^, and 100-500 μM (bulky) ATPγS for labeling. Lysate labeling was performed at 30°C for 30 minutes. Reactions were stopped by adding EDTA (pH 8) to a final concentration of 20 mM. PNBM was added to a final concentration of 2.5 mM and incubated for 1h at room temperature to alkylate the thiophosphorylated proteins in the labeled lysate. To stop alkylation, Laemmli buffer was added to final 1X and boiled for five minutes.

### Immunostaining

Cultured mESCs were washed with PBS (Gibco, #14190144), partially fixed in 4% formaldehyde (Thermo Scientific, #28908) in PBS for 3 minutes, washed with PBS, and covered with a permeabilizing and labeling reaction buffer (20 mM HEPES pH 7.5, 100 mM potassium acetate, 5 mM sodium acetate, 2 mM magnesium acetate, 1 mM EGTA (pH 8), 10 mM MgCl_2_, 5 mM MnCl_2_, 0.5 mM TCEP, 45 μg/ml digitonin (Sigma-Aldrich, #D141), 5 mM GTP, 0.2 mM ATP, 150 μM bulky ATP-γ-S, 1X phosphatase inhibitors (Roche, #4693132001)). Only a minimal amount of buffer (35-65 μl/cm^2^) was overlaid onto cells to cover the entire surface of the dish. Dishes were incubated at 30°C for 30 min with gentle shaking on a nutating mixer to label CDK2 substrates. After incubation, labeling was stopped and cells were fixed by aspirating labeling mixture and incubating dishes with a 20 mM EDTA + 4% formaldehyde in PBS solution for 10 min at RT on a nutating mixer. After a PBS rinse, labeled substrates were then alkylated with a 1 mM PNBM and 45 μg/ml digitonin solution in water for 30 min at RT on a nutating mixer. To quench the PNBM, the alkylation mixture was aspirated and 10 mM DTT in TBS was added to dishes, and finally rinsed with TBS.

Fixed and labeled cells were blocked with 5% BSA in TBS-T (0.1% Triton X-100) for 30 min at RT with gentle agitation and then incubated in 1:2000 rabbit anti-Thiophosphate ester antibody [51-8] (Abcam, #ab92570) solution containing 2% BSA in TBS-T overnight at 4°C overnight. Dishes were washed with TBS-T and incubated in 2% BSA secondary antibody solution containing 1:1000 donkey anti-rabbit IgG (H+L) Alexa Fluor™ 488 (Invitrogen, #A21206) and 500 ng/ml DAPI (Invitrogen, #D1306). Following another TBS-T wash, the plates were stored under light protection in PBS and imaged.

Phase contrast and immunofluorescence (DAPI and GFP channels) images were captured using the Lionheart FX system (Agilent BioTek) with Gen5 software (3.14). Images were processed in ImageJ^88^ to subtract background, then segmented and quantified using CellProfiler^89^.

### In situ thiophosphorylation and mass spectrometry

To process in situ labeled cells for lysis and thiophosphorylated protein enrichment, modifications were made to the in situ thiophosphorylation protocol above. Each cell line was grown in a 10 cm^2^ dish. The partial fixation step was omitted and 2 ml of labeling reaction buffer was directly added to the cells (20 mM HEPES pH 7.5, 100 mM potassium acetate, 5 mM sodium acetate, 2 mM magnesium acetate, 1 mM EGTA (pH 8), 10 mM MgCl_2_, 5 mM MnCl_2_, 0.5 mM TCEP, 45 μg/ml digitonin, 5 mM GTP, 0.2 mM ATP, 150 μM 6-Fu ATP-γ-S, 1X phosphatase inhibitor). Following incubation, labeling was terminated by adding EDTA (pH 8.0) to a final concentration of 20 mM. PNBM was then added to a final concentration of 1 mM and incubated for 2h at RT with gentle agitation on a nutating mixer to alkylate labeled substrates. Alkylation was quenched by adding DTT to a final concentration of 10 mM. Cells were subsequently scraped in the quenched labeling/alkylation mixture, transferred to tubes, and snap-frozen in liquid nitrogen until further processing.

Labeled lysates from three independent labeling experiments were thawed on ice and lysed in 1× RIPA buffer (Millipore, #20-188) bringing the final volume to 2.5 ml. For the following steps, all samples were kept on ice. To ensure complete lysis, samples were sonicated three times for 10s each at power setting 10 using a Microson Ultrasonic Cell Disruptor XL (Heat Systems Inc). Lysates were then transferred to pre-equilibrated PD-10 desalting columns (Cytiva, #17085101) and processed according to the gravity-flow protocol. Proteins were eluted in 3.5 ml of 1× RIPA with the first 1ml discarded, the subsequent 2.5 mL was collected and passed through a second pre-equilibrated PD-10 column. Final elution was performed in 3.5 mL of 1× RIPA and then supplemented with 1× protease inhibitor (Roche, #5056489001) and 0.1% SDS (UltraPure™ 10% SDS, Invitrogen, #15553-027). Protein concentrations were measured with Pierce™ BCA Protein Assay Kit (Thermo Scientific, #23225) in duplicate wells, with 25 µL of sample and 200 µL of working reagent per well. Total protein yields were consistent between samples (1.25-1.34 µg). Approximately 1% of each elution was reserved as an input sample for further analysis, and mixed with 4× Laemmli buffer (Bio-Rad, #1610747) containing 80 mM DTT, boiled at 95 °C for 5 min, centrifuged at 16,000 × g for 1 min, and stored at −20 °C. For immunoprecipitation, 10 µg of anti-thiophosphate ester antibody (Abcam, #ab133473) was added to the eluted proteins and incubated overnight at 4°C.

The following day, 150 µl of Dynabeads™ Protein G (Invitrogen, #10004D) was used per immunoprecipitation (IP). The beads were washed and resuspended in 1× RIPA-0.1% SDS solution, then added to lysate-antibody mixtures and incubated at 4 °C for 3.5h. After magnetic collection of the beads, but prior to washing, a 1% aliquot of the supernatant was saved as the unbound protein fraction and processed in the same manner as the input samples. Dynabeads were transferred to a new 1.5 ml Protein LoBind Tube (Eppendorf, #0030108442) during the first wash and subsequently washed five more times with 1× RIPA-0.1% SDS solution. From the sixth wash, approximately 5% aliquot was collected as an eluate sample. Proteins in this aliquot were mixed with 1× Laemmli buffer containing 20 mM DTT, boiled at 95 °C for 5 min, centrifuged at 16,000 × g for 1 min, and the supernatant was transferred to new tubes and stored at −20 °C. After removal of the sixth wash, Dynabeads were snap-frozen in liquid nitrogen and stored at −80 °C. Prior to mass-spectrometry, the input, unbound, and eluate samples were analyzed with Western blot using anti-Thiophosphate ester antibody to confirm substrate labeling and protein enrichment in the eluate fraction (data not shown).

Samples were solution digested with trypsin using S traps (Protifi), following the manufacturer’s instructions. Following lyophilization, dried peptides were resuspended in 5% acetonitrile, 0.05% TFA in water for mass spectrometry analysis on an Orbitrap Exploris 480 (Thermo Scientific) mass spectrometer. The peptides were separated on a 75 µm x 25 cm, 3 µm Acclaim PepMap reverse phase column (Thermo Scientific) at 300 nL/min using an UltiMate 3000 RSLCnano HPLC (Thermo Scientific) and eluted directly into the mass spectrometer. Parent full-scan mass spectra collected in the Orbitrap mass analyzer set to acquire data at 120,000 FWHM resolution and HCD fragment ions detected in the orbitrap at 15,000 resolution. Proteome Discoverer 3.0 (Thermo Scientific) was used to search the data against the mouse database from Uniprot using SequestHT. The search was limited to tryptic peptides, with maximally two missed cleavages allowed. Cysteine carbamidomethylation was set as a fixed modification, with methionine oxidation as a variable modification. The precursor mass tolerance was 10 ppm, and the fragment mass tolerance was 0.02 Da. The Percolator node was used to score and rank peptide matches using a 1% false discovery rate. Label-free quantitation (LFQ) of extracted ion chromatograms from MS1 spectra was performed using the Minora node in Proteome Discoverer.

### Mass-spectrometry data analysis

LFQ values were imported to R (v4.5.1) for further analysis. Missing values were imputed as 0.1% quantile LFQ of the respective sample. Data were log-transformed, and differential abundance between analog-sensitive CDK2 and wild-type CDK2 samples was calculated using linear models, followed by type II ANOVA (*car* package, v3.1-3) for statistical analysis. Proteins with positive coefficients and nominal p < 0.05 were classified as CDK2 substrates. Data were visualized as a volcano plot generated with *EnhancedVolcano* package (v1.26.0) and *ggplot2* (v3.5.2).

Gene Ontology enrichment analysis was performed using *clusterProfiler* (v4.16.0) using mouse annotations from *org.Mm.eg.db* (v3.21.0). All ontology domains were tested (ont = “all”) and pairwise term similarities were computed using pairwise_termsim command from enrichplot package (v1.28.2). Redundant categories were simplified based on similarity using Jaccard similarity coefficient with a cutoff of 0.7, keeping the categories with the lowest p-value. Enrichment results were then filtered to only keep categories with at least 3 genes and adjusted p-value < 0.05, and visualized with dotplot.

### Protein purification

Human cyclin-CDK fusion complexes were purified from *S. cerevisiae* as previously described^90^. Briefly, pRS425-prGAL1-3xFLAG-cyclin-L-CDK plasmids were transformed into yeast, grown in synthetic complete (SC) - leucine media with 2% raffinose, and expression was induced at OD 0.6-0.8 by adding 2% galactose, followed by incubation at 30 °C for 3h. The enzymes were purified using anti-FLAG M2 affinity agarose beads (Sigma-Aldrich, #A2220) and eluted with elution buffer (50 mM HEPES-KOH pH 7.6, 0.25 M KCl, 1 mM MgCl_2_, 1 mM EGTA, 5% Glycerol, 0.2 mg/mL 3×FLAG peptide (Sigma-Aldrich, #F4799)). Purified cyclin-CDK complexes were verified and required volumes were calculated relative to reference cyclin-CDK complexes after Western blot using anti-FLAG antibody (1:3000-5000, Sigma-Aldrich, #F1804) and donkey anti-mouse 800CW secondary antibody (1:12,000, LiCOR, #926-32212).

For substrate purifications, pET28a-6xHis plasmids were transformed into *E.coli* BL21(DE3)RIL cells (Agilent, #230245). Bacteria were grown in 2x YT media and protein expression was induced at OD 0.6 by adding 0.1-1 mM IPTG, and cultures were grown overnight at 18°C. Bacterial cells were collected and frozen in liquid nitrogen. Lysis was carried out using B-PER™ Complete Bacterial Protein Extraction Reagent (Thermo Scientific, #89822) supplemented with protease inhibitors (1 µg/ml pepstatin A, 1x protease inhibitor cocktail (1 µg/ml bestatin, 1 µg/ml leupeptin, 1 mM benzamidine HCl), 0.1 mM AEBSF, 0.6 µg/ml aprotinin). Lysates were clarified by centrifugation at 27,000g for 20 min at 4°C. His-tagged proteins were purified by gravity flow using nickel resin (HisPur™ Ni-NTA resin, Thermo Scientific, #88221). To increase binding specificity, 50 mM imidazole was added to the clarified lysate prior to loading onto the resin. The flowthrough was reapplied to the same resin to increase yield. Next, the resin was washed first with 20× resin volume and then 10× resin volume of 50 mM imidazole in the wash buffer (25 mM HEPES pH 7.4, 300 mM NaCl, 10% glycerol). Substrates were eluted in multiple steps: 3 times with 200 mM imidazole in the wash buffer, followed by 2 times with 500 mM imidazole in the wash buffer.

To purify GST-tagged C-terminus of HMCES, pGEX-GST-HMCES (271-353) plasmid was transformed into *E. coli* BL21(DE3)RIL bacteria (Agilent, #230245) and expression was induced with 0.1 mM IPTG overnight at 18 °C. Bacteria were lysed in B-PER™ Complete Bacterial Protein Extraction Reagent (Thermo Scientific #89822) containing protease inhibitors and sonicated with Microson ultrasonic cell disruptor XL (Heat Systems Inc) at power 10 for 15 seconds. GST-tagged proteins were batch bound to Pierce™ Glutathione Agarose beads (Thermo Scientific, #16101) at 4 °C for 2h, transferred to columns, and washed with 10× and 5× resin volume with 50 mM Tris pH 8.0, 100 mM potassium acetate, 25 mM magnesium acetate, and 0.1% Tween-20. 5 mM ATP was added to the second wash. The substrate was eluted 4 times with 50 mM Tris pH 8.0, 100 mM potassium acetate, 25 mM magnesium acetate, 10% glycerol, 15 mM glutathione.

Concentrations of all purified substrate preparations were measured against BSA standards. For that, substrate and BSA samples were loaded to 10% SDS-PAGE, stained with Coomassie (QC Colloidal Coomassie Stain, BioRad, #1610803), imaged with Typhoon FLA9500 (GE Healthcare Life Sciences), and quantified using ImageQuant TL software (v10.2-499). Purified HMCES preparations were also verified by Western blot using anti-HMCES antibody (1:2000, Sigma, #HPA044968), donkey anti-rabbit 800CW secondary antibody (1:10,000, LiCOR #926-32213), and imaged with Amersham Typhoon (Cytiva).

### In vitro kinase assays

In vitro kinase assays were performed as described in Zhang et al., 2025^91^ with minor modifications. Reactions contained 50 mM HEPES pH 7.4, 150 mM NaCl, 5 mM MgCl□, 0.5 mM DTT, 0.08 mg/mL BSA, 0.5 mM ATP-γ-S, and FLAG elution buffer to normalize reaction volumes across different cyclin-L-CDK complexes. Throughout the kinase reactions, His-HMCES variants were used at low micromolar concentrations (0.2-0.4 µM), while enzyme concentrations were maintained in the low nanomolar range. Histone H1 (Millipore, #14-155), a model substrate, was included in the CDK panel experiments at 10 µM concentration. Kinase reactions were incubated at room temperature for 5 or 8 min, then quenched with 20 mM EDTA and alkylated with 2.5 mM PNBM for 1 h at room temperature. Alkylation was terminated by addition of 20 mM DTT and 1× Laemmli buffer (Bio-Rad #1610747), followed by heating at 70 °C for 10 min and centrifugation at 16,000 × g for 1 min.

Samples were analyzed by SDS-PAGE followed by Western blotting. Total protein was visualized using the Revert™ 700 Total Protein Stain kit (LiCOR, #926-11010). The following antibodies were used: primary antibodies rabbit anti-Thiophosphate ester (1:2000, Abcam, #ab133473) and mouse anti-FLAG (1:3000-5000, Sigma-Aldrich, #F1804), and secondary antibodies donkey anti-rabbit 800CW and donkey anti-mouse 680CW antibody (1:12,000, LiCOR #926-32213 and #926-68022, respectively). Fluorescent signal was detected using Amersham Typhoon (Cytiva) and quantified with the ImageQuant TL software (v10.2-499).

In kinase assays with competitive peptides, full-length His-HMCES, C-terminal fragment of HMCES (GST-HMCES 271-353), or GST-H1 as a model substrate were used and incubated with cyclin E1-CDK2 for 5 min. Used peptides were as follows: p21 peptide containing RxL motif (SKACRRLFGPVDS, Peptide 2.0 Inc), peptide containing reverse RxL motif (rRxL, GSDKDFVIVRRPKLNRE, Peptide 2.0 Inc), and a peptide with mutated reverse RxL motif (rRxL mut, GSDKDFVAVRRPKLNRE, Peptide 2.0 Inc). All peptides were diluted in 50 mM HEPES pH 7.4 at 12 mM concentration and used at 500 µM concentration. Peptides were added along with substrate, before adding the enzyme and 5 min incubation. In reactions with no competitive peptide, equal volume of water was added to the reaction.

To determine the dynamics of HMCES phosphorylation, kinase assay with full-length His-HMCES and cyclin E1-CDK2 were performed in larger volume, and 10% aliquots were taken at each timepoint (0.5, 2.5, 5, 10, 20, 40, 60, 90, and 120 min). Reactions were stopped with a 2× Laemmli buffer containing 50 mM DTT. With site-mutated HMCES substrates, 20% aliquots were taken at different timepoints (0.5, 5, 20, 60 min). All samples were separated on 10% acrylamide, 50 µM Phos-tag, and 0.1 mM MnCl_2_-containing gels and visualized using anti-HMCES antibody (1:2000, Sigma, #HPA044968), donkey anti-rabbit 800CW antibody (1:12,000, LiCOR #926-32213) and imaged on Amersham Typhoon (Cytiva).

### In vitro coimmunoprecipitation and GST pull-down assays

75 fmoles of His-HMCES fragments, 1 µg of HMCES antibody (HPA044968-100UL), and 15 µl prewashed Dynabeads™ Protein G (Invitrogen, #10004D) per IP were mixed in co-IP buffer A (20 mM HEPES, pH 7.6, 150 mM NaCl, 10% glycerol, 0.5% NP-40, 1× protease inhibitor (Roche, #5056489001), and 0.2 µg/µl BSA) and incubated for 1h at 4 °C. Next, the beads were collected, supernatant was removed as much as possible, and the beads were washed 4 times with co-IP buffer A to remove unbound purified HMCES. Finally, the beads were resuspended in co-IP buffer A containing 1.875 fmoles of purified cyclin E1-CDK2, 1× protease inhibitor (Roche, #5056489001), and 0.2 µg/µl BSA. Immediately after resuspension, 2.5% of the sample was taken as an input sample. Next, the mixture was rotated 1h at 4 °C. After incubation, the beads were collected and 2.5% aliquot was taken from supernatant as an unbound sample. The beads were washed 4 times with co-IP buffer A and finally eluted in 2x Laemmli containing 100 mM DTT. The eluates were heated at 95 °C 5 min, collected by centrifugation at 16 000g for 1 min, and transferred to a new tube.

GST-HMCES C-terminus pulldown was performed with minor modifications. To remove glutathione, an aliquot of GST-HMCES preparation was dialyzed in 50 mM Tris pH 8.0, 100 mM potassium acetate, 25 mM magnesium acetate, and 10% glycerol solution at 4 °C for 1h, overnight, and 1h after changing the solution. Concentration of GST-HMCES was determined with Nanodrop measuring absorbance at 280 nm.

15 fmole of GST-HMCES_271-353_ was diluted in co-IP buffer B (50 mM HEPES, pH 7.6, 300 mM NaCl, 10% glycerol, 0.5% Triton X-100, 1x protease inhibitor (Roche, #5056489001), and 0.2 µg/µl BSA). For GST-HMCES binding, 20 µl of Pierce™ glutathione agarose slurry (Thermo Scientific, #16101) was taken for IP and incubated with GST-HMCES for 1.5h at 4 °C. Next, the resin was collected by centrifugation for 2 min at 700 g at 4 °C and supernatant was removed as much as possible. Next, the resin was resuspended in co-IP buffer C (20 mM HEPES pH 7.6, 200 mM NaCl, 1 mM EDTA, 0.5% Nonidet P-40, 10% glycerol, 1× protease inhibitor (Roche, #5056489001), 0.2 µg/µl BSA) with 1.875 fmole of cyclin E1-CDK2 and rotated at 4 °C for 3.5h. 5% of input sample was taken before moving the mixture to 4 °C. After incubation, the resin was collected again by centrifugation, and 5% aliquot was taken from supernatant as unbound sample. The resin was washed 4 times with co-IP buffer C for 5-10 min at 4 °C and collected each time with centrifugation. Before elution, resin was centrifuged again, and the wash solution was removed thoroughly. Bound proteins were eluted in 2× Laemmli containing 100 mM DTT, heated at 95 °C 5 min, collected by centrifugation at 16 000g for 1 min, and transferred to a new tube. GST-HMCES_271-353_ pulldown experiment was repeated 2 times.

All samples were separated using in-house made 4-20% gradient gels and analyzed by Western blotting with anti-HMCES and anti-FLAG antibodies or using Revert™ 700 Total Protein Stain kit (LiCOR, #926-11010) to detect total GST-HMCES.

### Fluorescence polarization assay

Equal amounts of His-HMCES WT and triple alanine-mutated HMCES (named 3A) were taken to kinase assay with cyclin E1-CDK2 and phosphorylated with regular ATP for 1h at room temperature. To prevent the reaction (inactive cyclin E1-CDK2 sample) or stop the reaction after 1h, final concentration of 20 mM EDTA was added to the reaction. Immediately after adding cyclin E1-CDK2 to HMCES, and right before stopping the reaction, a 2% aliquot was taken for analysis on Phos-tag gel.

DNA binding was determined as described in Halabelian et al., 2019^70^. For that, each kinase assay sample was divided to three wells (10 ul per well) on black polystyrene 96-well plate (Corning, #3686), and 40 µl of DNA binding buffer (20□mM HEPES pH 7.4, 140□mM KCl, 5□mM NaCl, 0.1□mM EDTA, 0.01% Triton X-100, 0.2□mM TCEP (Thermo Scientific, #77720)), and 10 nM DNA probe was added. 6-FAM-5’-TCTTCTGGTCCGGATGGTAGTTAAGTGTTGAG-3’ (Integrated DNA Technologies) ssDNA sequence was used as the probe according to Halabelian 2019^70^. The binding reaction was incubated at room temperature for 30 min and light polarization was measured with Agilent BioTek Synergy Neo2 Microplate Reader.

### Electrophoretic mobility shift assay (EMSA)

Equal amounts of full-length His-HMCES or His-HMCES_1-270_ were used as starting material and serially diluted two-fold, after which 1 µl was added to EMSA reactions. To determine total free oligo, EMSA reactions were performed with the elution buffer from protein purification step instead of HMCES.

To test the effect of phosphorylation, equal amounts of His-HMCES WT and triple alanine-mutated HMCES (20 µM in kinase reaction) were phosphorylated with regular ATP by cyclin E1-CDK2 for 1h at room temperature. To obtain different concentrations of HMCES, kinase assay reactions were sequentially diluted 2-fold in the kinase assay reaction buffer and 1 µl was added to EMSA reactions. Successful phosphorylation of HMCES was later confirmed on Phos-tag gel and equal total levels were confirmed using regular SDS-PAGE.

EMSA reactions were performed in total 5 µl, containing 1× binding buffer (20 mM HEPES pH 7.4, 140 mM KCl, 5 mM NaCl, 0.1 mM EDTA, 0.01% Triton X-100, 2% Ficoll), 0.2 µg/µl BSA, 0.2 mM TCEP (Thermo Scientific, #77720), and 16.7 nM oligo 6-FAM-5’-TCTTCTGGTCCGGATGGTAGTTAAGTGTTGAG-3’ (Integrated DNA Technologies). Binding reactions were performed for 30 min in the dark at RT.

EMSA gels (10% acrylamide, 0.25× TBE, 0.01% NP-40) were pre-run in 0.5× TBE for 20-30 min at 100V. All the 5 µl of EMSA reactions were loaded to the gel and ran in dark for 1h at 100 V at RT. Finally, the gels were imaged using Typhoon FLA9500 (GE Healthcare Life Sciences).

For analysis, band intensities for free oligonucleotide were quantified in ImageQuant TL (v10.2-499). Bound oligonucleotide was calculated indirectly by subtracting the free oligonucleotide signal measured in each sample lane from that measured in the elution buffer only control. All data were analyzed and plotted using GraphPad Prism (v10.1.1). Data were first min-max normalized and then fit by nonlinear regression using the one site-specific binding model to generate binding curves and calculate K_d_ values.

### AP site analysis and viability assay

ESCs were plated at 25,000 cells/well in gelatin-coated 12-well dishes in ESGRO-2i complete medium (Sigma-Aldrich, #SF016). The next day, cells were treated with doxycycline (100 ng/mL) to induce exogenous siRNA-resistant HMCES expression and transfected overnight with Lipofectamine RNAiMAX (Invitrogen, #13778150) and 15 pmol of siGENOME Non-Targeting siRNA Pool #1 (Horizon Discovery, #D-001206-13-05) or siGENOME Mouse Hmces (232210) siRNA SMARTpool (Horizon Discovery, #M-062972-00-0005). The following morning, transfected cells were fed with fresh media containing doxycycline (100 ng/mL) and incubated for 24h. The next day, cells were treated with 10 mM MMS and returned to the 37°C incubator for 1h. After 1h, the media was removed, cells were rinsed with PBS, media was refreshed, and cells were returned to the 37°C incubator for 2h.

To measure AP sites, cells were lysed and genomic DNA was extracted with a Quick-DNA MicroPrep kit (Zymo, #D3020). Genomic DNA concentration was measured with a spectrophotometer, and 1 µg of genomic DNA was labeled with a biotinylated Aldehyde Reactive Probe (ARP) using the Nucleostain DNA Damage Quantification Kit (Dojindo Laboratories, #DK02). The ARP-labeled DNA was then purified following manufacturer’s instructions. Purified ARP-labeled genomic DNA concentration was measured with a Qubit dsDNA HS Assay Kit (Q32851) on a Qubit 4 Fluorometer. For AP site measurement assays, 202.5 ng of purified ARP-labeled genomic DNA was resuspended in 200 µL of TE. Experimental samples and ARP-DNA standard solutions were aliquoted onto 96-well plates and AP sites were measured based on manufacturer’s instructions. Optical density at 650 nm was measured using Biotek Synergy plate reader.

### Cell viability assay

15,000 cells/well of mESCs were seeded in 24-well gelatin-coated plates in ESGRO-2i complete medium (Sigma-Aldrich, #SF016). The following day, doxycycline treatment (100 ng/mL) and siRNA transfections were performed using Lipofectamine RNAiMAX (Thermo Fisher Scientific, # 13778150) according to the manufacturer’s instructions. The next day, the cell media was replaced with a fresh dox-containing growth media for another 24h. The following day, cells were treated with 2 mM MMS or 1 mM KBrO□, as determined by IC_50_ (data not shown). Cell viability was assessed by Trypan Blue exclusion using Countess 3 Automated Cell Counter (Thermo Fisher Scientific; software release 1.4.1625.619). Briefly, cells were detached with Accutase (Gibco, #00-4555-56), pelleted, and resuspended in DMEM/F12 (Gibco, #11039-021). Cell suspensions were individually mixed 1:1 (v/v) with 0.4% Trypan Blue solution (Thermo Fisher Scientific, #T10282), loaded onto a Countess chamber slide, and measured. Relative cell count was calculated as the percentage of viable cells in each treated condition relative to the untreated control.

### Statistical analysis and graphical representation of the data

For cyclin-CDK panel experiments (Fig. 3b,c), quantification of thiophosphorylated and total protein levels was performed using ImageQuant TL software (v10.2-499). Relative substrate phosphorylation was calculated as the ratio of thiophosphorylated signal to total substrate signal. HMCES phosphorylation was then further normalized to Histone H1 phosphorylation levels. Data from three independent experiments (n = 3) were log-transformed, mean-centered, and auto-scaled prior to statistical analysis. Mean values and standard errors of the mean (SEM) were calculated on auto-scaled data. For graphical presentation, mean and mean ± SEM were back-transformed to the linear scale with error bars representing the upper and lower limits of the back-transformed mean ± SEM. Data are shown relative to the average HMCES phosphorylation by cyclin E1-CDK2. Statistical significance was assessed using one-way repeated-measures ANOVA with Holm-Šídák post hoc test on auto-scaled data in GraphPad Prism (v10.1.1).

For peptide competition experiments (Fig. 3g,h), relative phosphorylation of each substrate by wild-type cyclin E1-CDK2 in the absence of peptide was set to 1. Data were log-transformed prior to statistical analysis. Results are presented as mean ± SEM, with n = 3. Statistical analysis was performed using two-way repeated-measures ANOVA and Dunnett’s post hoc test on log-transformed data in GraphPad Prism (v10.1.1).

For experiments testing the effect of hydrophobic patch mutant (Fig. 3 i,j), thiophosphorylated and total substrate levels were quantified as described above. Relative phosphorylation by wild-type cyclin E1-CDK2 was set to 1. Data from three independent experiments (n = 3) were log-transformed and mean ± SEM were calculated. For visualization, values were back-transformed to linear scale with error bars indicating back-transformed mean ± SEM. Statistical analysis was performed using one-way repeated-measures ANOVA with Holm-Šídák post hoc test on log-transformed data in GraphPad Prism (v10.1.1).

For analyzing phosphorylation of HMCES fragments (Fig. 4a,b), data were log-transformed and shown relative to the phosphorylation of full-length HMCES. Statistical significance was evaluated using one-way repeated-measures ANOVA with mixed-effects models followed by Dunnett’s post hoc test on log-transformed data in GraphPad Prism (v10.1.1).

For HMCES phosphomutant experiments (Fig. 4e,f), data were log-transformed prior to calculation of mean and SEM. Sample sizes were n = 3 for cyclin E1-CDK2 and n = 2 for all other cyclin-CDK complexes. No statistical testing was performed for these experiments.

For polarization-based DNA-binding assays (Fig. 5f), measurements from three technical replicates (wells) were averaged per condition. Relative light polarization (ΔP) was calculated by subtracting the mean polarization signal from samples containing no protein. In total, four independent experiments were performed (n = 4). Statistical significance was assessed using one-way repeated-measures ANOVA with Holm-Šídák post hoc test in GraphPad Prism (v10.1.1).

For analyzing AP-site data (Fig. 5h), the number of AP sites was normalized to the AP-site level in corresponding non-treated cells transfected with siCTRL. The data was then log-transformed, and statistical analysis was carried out using two-way repeated measurements ANOVA with Holm-Šídák post hoc test in Graphpad Prism (v10.1.1).

For analyzing cell numbers (Fig. 5i), data were normalized to cell numbers in nontreated cells in respectively silenced and HMCES-expressing cells. Normalized data was log-transformed, and statistical significance was calculated using two-way repeated measurement ANOVA with Holm-Šídák post hoc test in Graphpad Prism (v10.1.1).

## Acknowledgements

We would like to thank all members of the Shariati and Koivomagi lab for insightful discussions and for sharing reagents. This research was funded by support from National Institutes of Health (NIH) under award number K12GM139185 and the Institute for the Biology of Stem Cells (IBSC) at UC Santa Cruz (B.R.T), the NIH/NIGMS through a Pathway to Independence award under award number K99GM126027/R00GM126027 (S.A.S), Maximizing Investigator Award under award number R35GM147395 (S.A.S), a start-up package from the University of California (A.S), Santa Cruz (S.A.S.) and a seed grant from Genomics Institute of UC Santa Cruz (S.A.S), the Intramural Research Program of the National Institutes of Health (NIH Grant ZIA BC 012010-06 to T.H.S. and ZIA BC 012133 to M.K.). The contributions of the NIH author(s) are considered Works of the United States Government. The findings and conclusions presented in this paper are those of the author(s) and do not necessarily reflect the views of the NIH and the U.S. Department of Health and Human Services.

## Extended Data

**Extended Data Figure 1.**
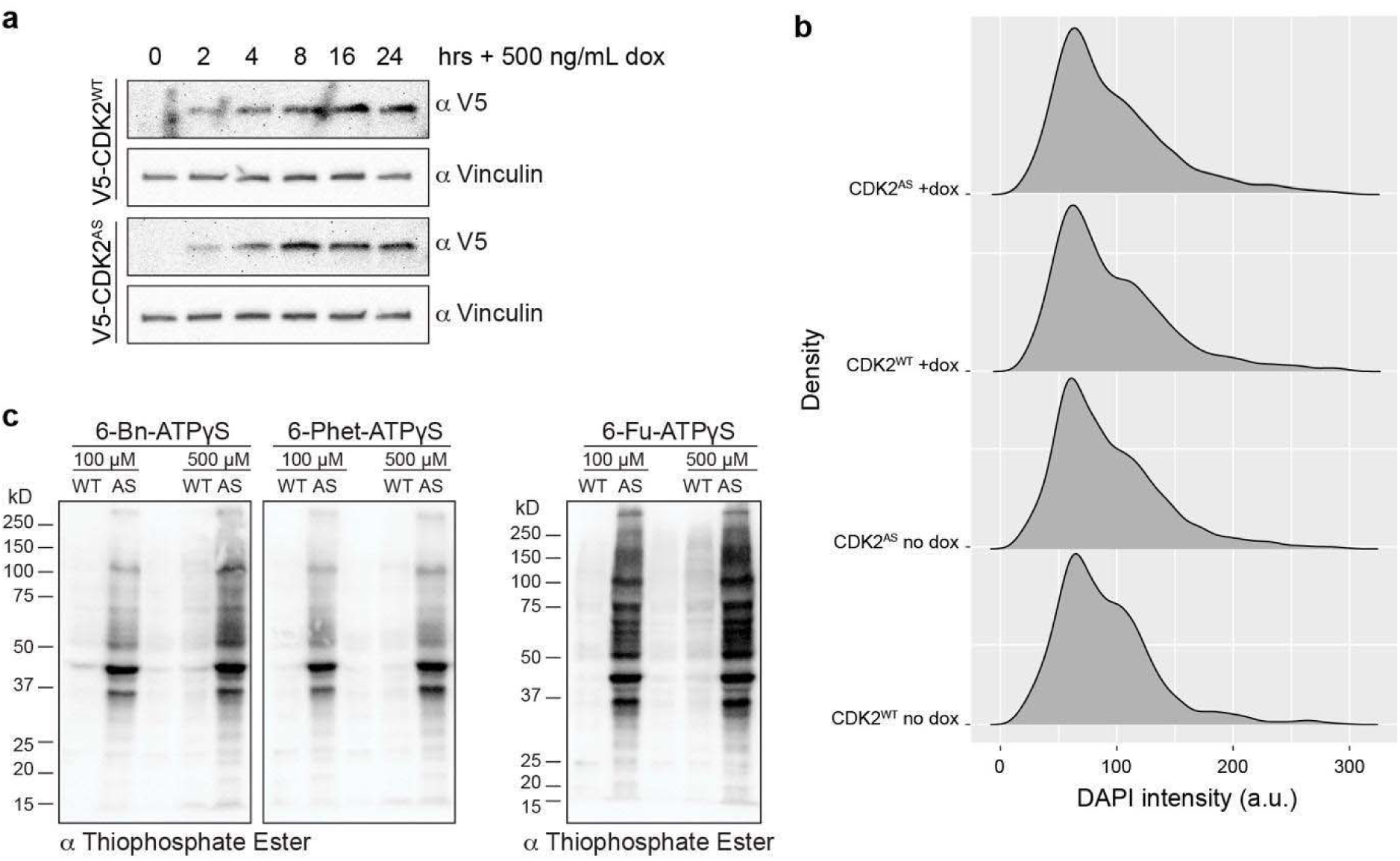
Comparison of transgenic CDK2^WT^ and CDK2^AS^ mESCs. **a**, Temporal dynamics of the dox-inducible V5-CDK2 expression system. Transgenic mESC lines were treated with 500 ng/mL dox, lysed, analyzed by SDS-PAGE using anti-V5 for exogenous CDK2 levels or anti-Vinculin antibody as loading control. **b**, Cell cycle profiles of transgenic CDK2^WT^ and CDK2^AS^ mESCs after 24h of 500 ng/mL dox induction. Cells were fixed, permeabilized, and DNA was stained with DAPI. Histograms are normalized to represent probability density, where the total area under each histogram equals one. **c**, In-cell labeling of CDK2 substrates using different bulky-ATP analogs at 100 μM or 500 μM concentrations.

**Extended Data Figure 2.**
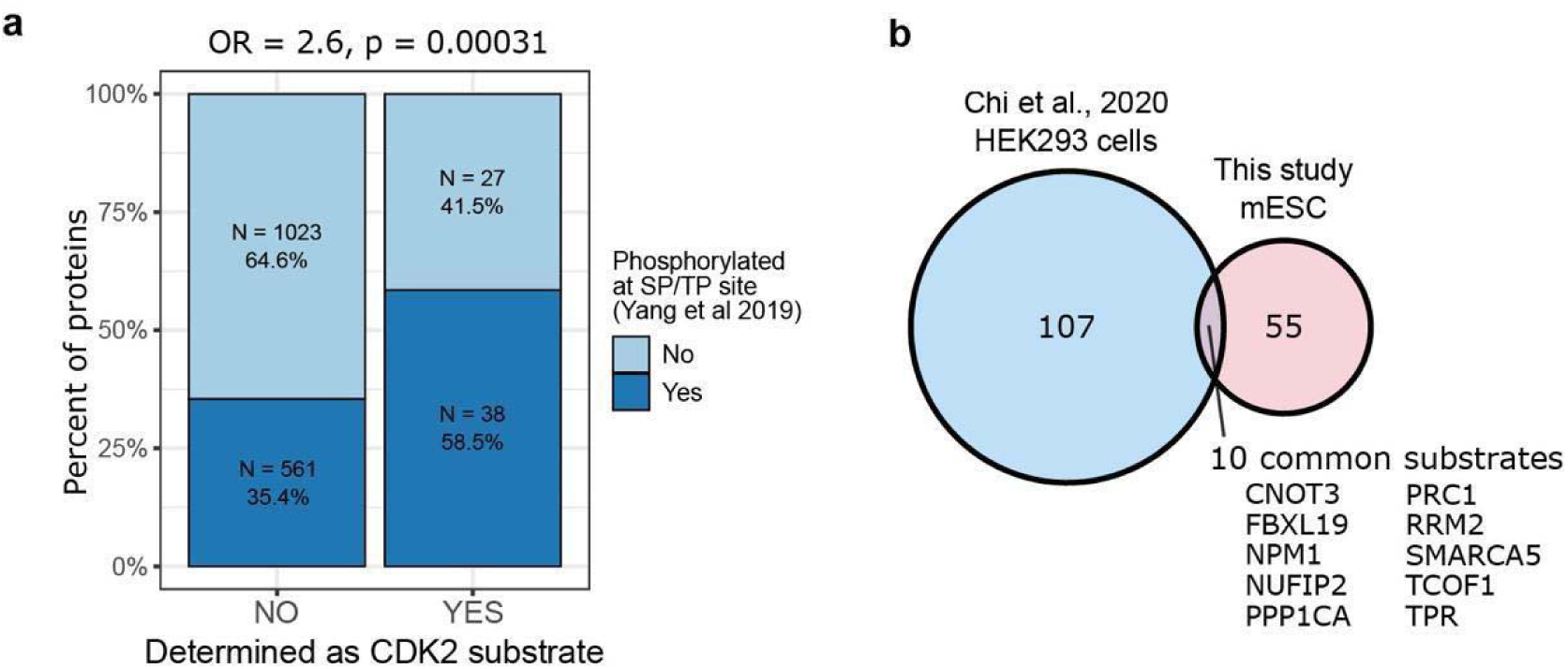
Comparison of CDK2 targets with previous studies. **a**, Overlap between the CDK2 substrates identified in our mass-spectrometric analysis and proteins previously reported by Yang et al., 2019^50^ to be phosphorylated at SP or TP sites in mESCs. Proteins determined not to be CDK2 substrates were used as a control group. **b**, Venn diagram comparing CDK2 substrates identified in this study with those reported by Chi et al., 2020^17^ in HEK293 cells.

**Extended Data Figure 3.**
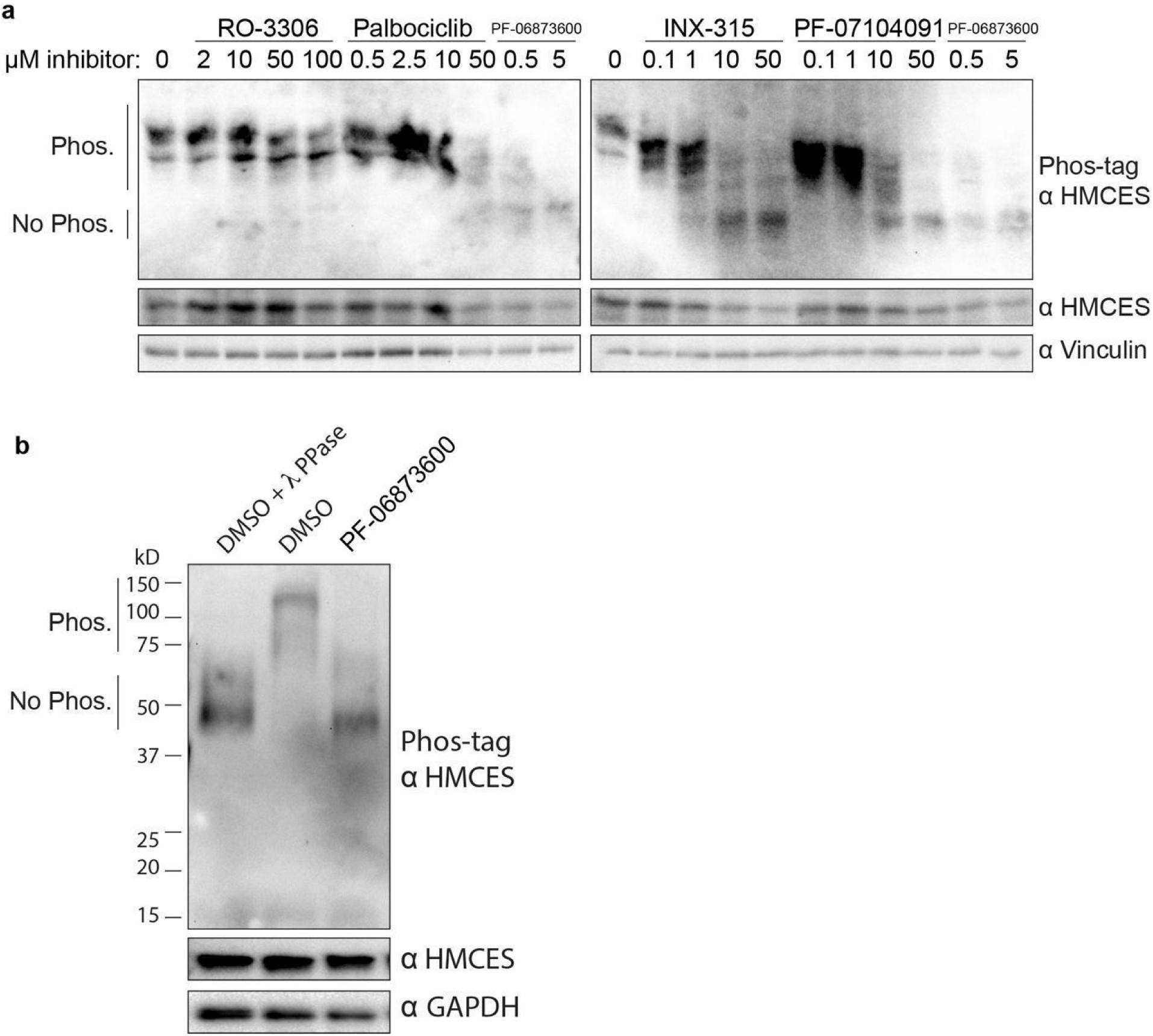
HMCES phosphorylation is CDK2-dependent in cells. **a**, mESCs were treated with the indicated CDK inhibitors at the indicated concentrations or DMSO for 1h, lysed, and analyzed by Phos-tag or regular SDS-PAGE using anti-HMCES antibody or anti-Vinculin as loading control. **b**, mESCs were treated with PF-06873600 or DMSO for 1h and lysed. The lysates treated with λ phosphatase where indicated and analyzed by Phos-tag or regular SDS-PAGE using anti-HMCES or anti-GAPDH antibody as loading control.

**Extended Data Figure 4.**
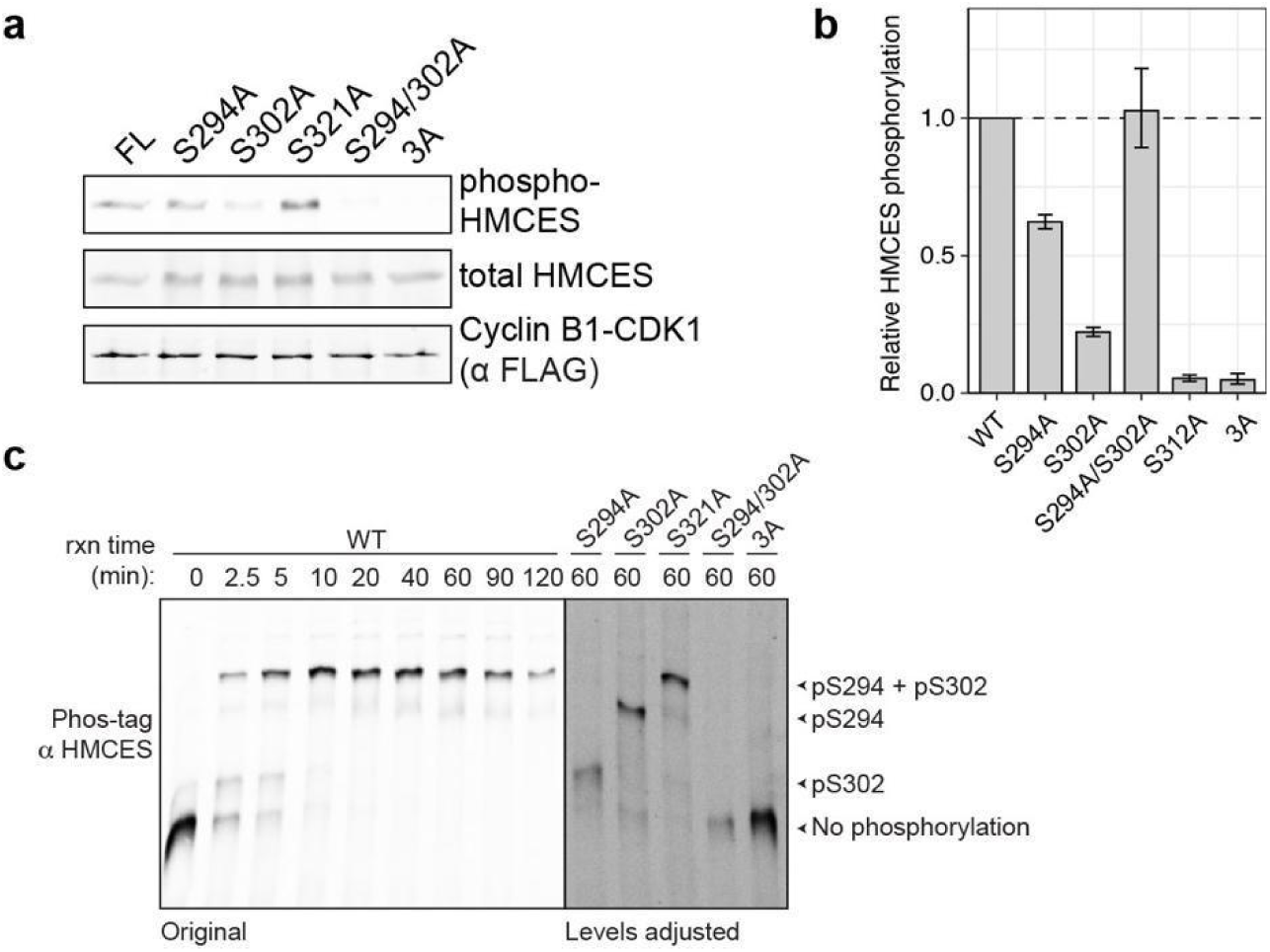
HMCES phosphorylation by cyclin B1-CDK1 and the dynamics of site-specific HMCES phosphorylation by cyclin E1-CDK2. **a**, In vitro kinase assay with full-length His-HMCES or HMCES phosphorylation variants incubated with cyclin B1-CDK1 for 5 min. **b**, Quantification of kinase assays shown on panel **a**. Data is shown relative to wild-type HMCES phosphorylation by each cyclin-CDK complex (mean ± SEM, n = 2). **c**, In vitro kinase assay with wild-type HMCES (HMCES WT) and cyclin E1-CDK2 for the indicated time course and separated by Phos-tag using anti-HMCES antibody along with phosphosite mutants for comparison. Note that this blot is a lower exposure of the blot in Fig. 4a.

**Extended Data Figure 5.**
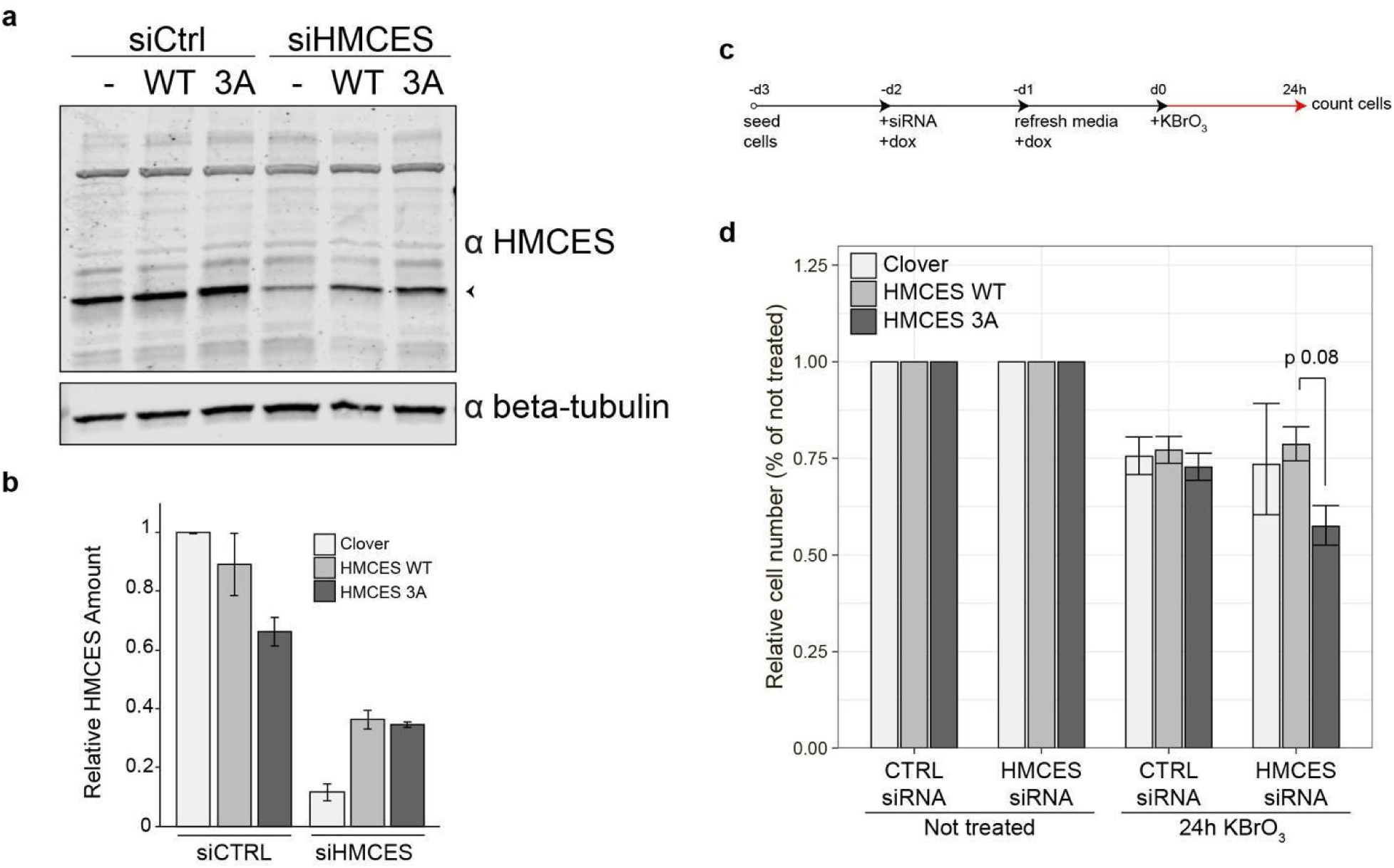
HMCES phosphorylation promotes cell viability upon KBrO_3_ treatment. **a**, Characterization of endogenous HMCES depletion and exogenous HMCES expression in mESCs. mESCs were treated with siRNAs against non-targeting control (siCTRL) or HMCES (siHMCES) in the presence of 100 ng/mL dox to induce expression of Clover protein (denoted as -), siRNA-resistant HMCES WT, or siRNA-resistant HMCES 3A for 48h. Media was refreshed with dox after 24h. Cells were lysed and analyzed by SDS-PAGE using anti-HMCES or anti-beta-tubulin antibody as loading control. **b**, Quantification of immunoblots shown on panel **a**. **c**, Schematic of cell viability assay experiment after KBrO_3_ treatment. Endogenous HMCES was depleted with siRNA and doxycycline-inducible Clover, siRNA-resistant wild-type HMCES (HMCES WT), or siRNA-resistant triple alanine-mutated HMCES (HMCES 3A) were induced in transgenic mESC cell lines. Cells were treated with 1 mM KBrO□ for 24h and then counted. **d**, Relative cell numbers after 24h of KBrO□. Data is shown relative to cell numbers in nontreated cells in respectively silenced and HMCES-expressing cells. (mean ± SEM, n = 5).

## Supplementary Information and Tables

**Supplementary Table 1. Analog-sensitive CDK2 mass spectrometry analysis and CDK site motif quantification.**

**Supplementary Table 2. Plasmids used in this study.**

## References

1. Davidson, K. C., Mason, E. A. & Pera, M. F. The pluripotent state in mouse and human. Development 142, 3090–3099 (2015).

2. Li, M. & Belmonte, J. C. I. Ground rules of the pluripotency gene regulatory network. Nat. Rev. Genet. 18, 180–191 (2017).

3. Ballabeni, A. et al. Cell cycle adaptations of embryonic stem cells. Proc. Natl. Acad. Sci. U. S. A. 108, 19252–19257 (2011).

4. Liu, L., Michowski, W., Kolodziejczyk, A. & Sicinski, P. The cell cycle in stem cell proliferation, pluripotency and differentiation. Nat. Cell Biol. 21, 1060–1067 (2019).

5. Coronado, D. et al. A short G1 phase is an intrinsic determinant of naïve embryonic stem cell pluripotency. Stem Cell Res. 10, 118–131 (2013).

6. Boward, B., Wu, T. & Dalton, S. Concise review: Control of cell fate through cell cycle and pluripotency networks. Stem Cells 34, 1427–1436 (2016).

7. Kareta, M. S., Sage, J. & Wernig, M. Crosstalk between stem cell and cell cycle machineries. Curr. Opin. Cell Biol. 37, 68–74 (2015).

8. DeVeale, B. et al. G1/S restriction point coordinates phasic gene expression and cell differentiation. Nat. Commun. 13, 3696 (2022).

9. Lange, C. & Calegari, F. Cdks and cyclins link G1 length and differentiation of embryonic, neural and hematopoietic stem cells. Cell Cycle 9, 1893–1900 (2010).

10. Rocha, C. R. R., Lerner, L. K., Okamoto, O. K., Marchetto, M. C. & Menck, C. F. M. The role of DNA repair in the pluripotency and differentiation of human stem cells. Mutat. Res. 752, 25–35 (2013).

11. Ter Huurne, M. & Stunnenberg, H. G. G1-phase progression in pluripotent stem cells. Cell. Mol. Life Sci. 78, 4507–4519 (2021).

12. Pauklin, S. & Vallier, L. The cell-cycle state of stem cells determines cell fate propensity. Cell 156, 1338 (2014).

13. Kareta, M. S. et al. Inhibition of pluripotency networks by the Rb tumor suppressor restricts reprogramming and tumorigenesis. Cell Stem Cell 16, 39–50 (2015).

14. Thompson, P. S. & Cortez, D. New insights into abasic site repair and tolerance. DNA Repair (Amst.) 90, 102866 (2020).

15. Mohni, K. N. et al. HMCES maintains genome integrity by shielding abasic sites in single-strand DNA. Cell 176, 144–153.e13 (2019).

16. Chi, Y. et al. Identification of CDK2 substrates in human cell lysates. Genome Biol. 9, R149 (2008).

17. Chi, Y. et al. A novel landscape of nuclear human CDK2 substrates revealed by in situ phosphorylation. Sci. Adv. 6, eaaz9899 (2020).

18. Elphick, L. M. et al. A quantitative comparison of wild-type and gatekeeper mutant cdk2 for chemical genetic studies with ATP analogues. Chembiochem 10, 1519–1526 (2009).

19. Wohlbold, L. et al. Chemical genetics reveals a specific requirement for Cdk2 activity in the DNA damage response and identifies Nbs1 as a Cdk2 substrate in human cells. PLoS Genet. 8, e1002935 (2012).

20. Hertz, N. T. et al. Chemical genetic approach for kinase-substrate mapping by covalent capture of thiophosphopeptides and analysis by mass spectrometry. Curr. Protoc. Chem. Biol. 2, 15–36 (2010).

21. Allen, J. J. et al. A semisynthetic epitope for kinase substrates. Nat. Methods 4, 511–516 (2007).

22. Ding, S. et al. Efficient transposition of the piggyBac (PB) transposon in mammalian cells and mice. Cell 122, 473–483 (2005).

23. Matsushime, H. et al. D-type cyclin-dependent kinase activity in mammalian cells. Mol. Cell. Biol. 14, 2066–2076 (1994).

24. Michowski, W. et al. Cdk1 controls global epigenetic landscape in embryonic stem cells. Mol. Cell 78, 459–476.e13 (2020).

25. Hein, J. B. & Nilsson, J. Interphase APC/C-Cdc20 inhibition by cyclin A2-Cdk2 ensures efficient mitotic entry. Nat. Commun. 7, 10975 (2016).

26. Ohtoshi, A., Maeda, T., Higashi, H., Ashizawa, S. & Hatakeyama, M. Human p55(CDC)/Cdc20 associates with cyclin A and is phosphorylated by the cyclin A-Cdk2 complex. Biochem. Biophys. Res. Commun. 268, 530–534 (2000).

27. Jiang, W. et al. PRC1: a human mitotic spindle-associated CDK substrate protein required for cytokinesis. Mol. Cell 2, 877–885 (1998).

28. Zhu, C., Lau, E., Schwarzenbacher, R., Bossy-Wetzel, E. & Jiang, W. Spatiotemporal control of spindle midzone formation by PRC1 in human cells. Proc. Natl. Acad. Sci. U. S. A. 103, 6196–6201 (2006).

29. Silva Cascales, H., et al. Cyclin A2 localises in the cytoplasm at the S/G2 transition to activate PLK1. Life Sci. Alliance 4, e202000980 (2021).

30. Tavernier, N. et al. Bora phosphorylation substitutes in trans for T-loop phosphorylation in Aurora A to promote mitotic entry. Nat. Commun. 12, 1899 (2021).

31. Okuda, M. et al. Nucleophosmin/B23 is a target of CDK2/cyclin E in centrosome duplication. Cell 103, 127–140 (2000).

32. Tarapore, P. et al. Thr199 phosphorylation targets nucleophosmin to nuclear speckles and represses pre-mRNA processing. FEBS Lett. 580, 399–409 (2006).

33. Tokuyama, Y., Horn, H. F., Kawamura, K., Tarapore, P. & Fukasawa, K. Specific phosphorylation of nucleophosmin on Thr(199) by cyclin-dependent kinase 2-cyclin E and its role in centrosome duplication. J. Biol. Chem. 276, 21529–21537 (2001).

34. Deng, M. et al. Identification and functional analysis of a novel cyclin e/cdk2 substrate ankrd17. J. Biol. Chem. 284, 7875–7888 (2009).

35. Gu, J. et al. Cell cycle-dependent regulation of a human DNA helicase that localizes in DNA damage foci. Mol. Biol. Cell 15, 3320–3332 (2004).

36. Spencer, S. L. et al. The proliferation-quiescence decision is controlled by a bifurcation in CDK2 activity at mitotic exit. Cell 155, 369–383 (2013).

37. Koundrioukoff, S. et al. A direct interaction between proliferating cell nuclear antigen (PCNA) and Cdk2 targets PCNA-interacting proteins for phosphorylation. J. Biol. Chem. 275, 22882–22887 (2000).

38. Boudrez, A., Beullens, M., Waelkens, E., Stalmans, W. & Bollen, M. Phosphorylation-dependent interaction between the splicing factors SAP155 and NIPP1. J. Biol. Chem. 277, 31834–31841 (2002).

39. Murthy, T. et al. Cyclin-dependent kinase 1 (CDK1) and CDK2 have opposing roles in regulating interactions of splicing factor 3B1 with chromatin. J. Biol. Chem. 293, 10220–10234 (2018).

40. Seghezzi, W. et al. Cyclin E associates with components of the pre-mRNA splicing machinery in mammalian cells. Mol. Cell. Biol. 18, 4526–4536 (1998).

41. Ramani, K. et al. S-adenosylmethionine inhibits la ribonucleoprotein domain family member 1 in murine liver and human liver cancer cells. Hepatology 75, 280–296 (2022).

42. Chung, K.-P. et al. Multi-kinase framework promotes proliferation and invasion of lung adenocarcinoma through activation of dynamin-related protein 1. Mol. Oncol. 15, 560–578 (2021).

43. Hu, G. et al. A genome-wide RNAi screen identifies a new transcriptional module required for self-renewal. Genes Dev. 23, 837–848 (2009).

44. Sarkar, M. et al. CNOT3 interacts with the Aurora B and MAPK/ERK kinases to promote survival of differentiating mesendodermal progenitor cells. Mol. Biol. Cell 32, ar40 (2021).

45. Zheng, X. et al. Cnot1, Cnot2, and Cnot3 maintain mouse and human ESC identity and inhibit extraembryonic differentiation. Stem Cells 30, 910–922 (2012).

46. Zheng, X. et al. CNOT3-dependent mRNA deadenylation safeguards the pluripotent state. Stem Cell Reports 7, 897–910 (2016).

47. Higashi, H. et al. Differences in substrate specificity between Cdk2-cyclin A and Cdk2-cyclin E in vitro. Biochem. Biophys. Res. Commun. 216, 520–525 (1995).

48. Johnson, J. L. et al. An atlas of substrate specificities for the human serine/threonine kinome. Nature 613, 759–766 (2023).

49. Kitagawa, M. et al. The consensus motif for phosphorylation by cyclin D1-Cdk4 is different from that for phosphorylation by cyclin A/E-Cdk2. EMBO J. 15, 7060–7069 (1996).

50. Yang, P. et al. Multi-omic profiling reveals dynamics of the phased progression of pluripotency. Cell Syst. 8, 427–445.e10 (2019).

51. Neganova, I. et al. An important role for CDK2 in G1 to S checkpoint activation and DNA damage response in human embryonic stem cells. Stem Cells 29, 651–659 (2011).

52. Peña-Gómez, M. J. et al. HMCES corrupts replication fork stability during base excision repair in homologous recombination-deficient cells. Sci. Adv. 11, eads3227 (2025).

53. Rua-Fernandez, J. et al. Self-reversal facilitates the resolution of HMCES DNA-protein crosslinks in cells. Cell Rep. 42, 113427 (2023).

54. Semlow, D. R., MacKrell, V. A. & Walter, J. C. The HMCES DNA-protein cross-link functions as an intermediate in DNA interstrand cross-link repair. Nat. Struct. Mol. Biol. 29, 451–462 (2022).

55. Srivastava, M. et al. HMCES safeguards replication from oxidative stress and ensures error-free repair. EMBO Rep. 21, e49123 (2020).

56. Thompson, P. S., Amidon, K. M., Mohni, K. N., Cortez, D. & Eichman, B. F. Protection of abasic sites during DNA replication by a stable thiazolidine protein-DNA cross-link. Nat. Struct. Mol. Biol. 26, 613–618 (2019).

57. Koledova, Z. et al. Cdk2 inhibition prolongs G1 phase progression in mouse embryonic stem cells. Stem Cells Dev. 19, 181–194 (2010).

58. Stead, E. et al. Pluripotent cell division cycles are driven by ectopic Cdk2, cyclin A/E and E2F activities. Oncogene 21, 8320–8333 (2002).

59. Ahuja, A. K. et al. A short G1 phase imposes constitutive replication stress and fork remodelling in mouse embryonic stem cells. Nat. Commun. 7, 10660 (2016).

60. Maynard, S. et al. Human embryonic stem cells have enhanced repair of multiple forms of DNA damage. Stem Cells 26, 2266–2274 (2008).

61. Tichy, E. D. et al. Mouse embryonic stem cells, but not somatic cells, predominantly use homologous recombination to repair double-strand DNA breaks. Stem Cells Dev. 19, 1699–1711 (2010).

62. Tichy, E. D. et al. The abundance of Rad51 protein in mouse embryonic stem cells is regulated at multiple levels. Stem Cell Res. 9, 124–134 (2012).

63. Arora, M. et al. Rapid adaptation to CDK2 inhibition exposes intrinsic cell-cycle plasticity. Cell 186, 2628–2643.e21 (2023).

64. Dietrich, C. et al. Correction: INX-315, a selective CDK2 inhibitor, induces cell cycle arrest and senescence in solid tumors. Cancer Discov. 15, 2185 (2025).

65. Topacio, B. R. et al. Cyclin D-Cdk4,6 drives cell-cycle progression via the retinoblastoma protein’s C-terminal helix. Mol. Cell 74, 758–770.e4 (2019).

66. Kelso, S. et al. Bipartite binding of the N terminus of Skp2 to cyclin A. Structure 29, 975–988.e5 (2021).

67. Iakoucheva, L. M. et al. The importance of intrinsic disorder for protein phosphorylation. Nucleic Acids Res. 32, 1037–1049 (2004).

68. Schulman, B. A., Lindstrom, D. L. & Harlow, E. Substrate recruitment to cyclin-dependent kinase 2 by a multipurpose docking site on cyclin A. Proc. Natl. Acad. Sci. U. S. A. 95, 10453–10458 (1998).

69. Örd, M. et al. High-throughput investigation of cyclin docking interactions reveals the complexity of motif binding determinants. Nat. Commun. 16, 7622 (2025).

70. Halabelian, L. et al. Structural basis of HMCES interactions with abasic DNA and multivalent substrate recognition. Nat. Struct. Mol. Biol. 26, 607–612 (2019).

71. Kurisu, S. et al. Quantitation of DNA damage by an aldehyde reactive probe (ARP). Nucleic Acids Res. Suppl. 1, 45–46 (2001).

72. Rubin, S. M., Sage, J. & Skotheim, J. M. Integrating old and new paradigms of G1/S control. Mol. Cell 80, 183–192 (2020).

73. Saykali, B. et al. Lineage-specific CDK activity dynamics characterize early mammalian development. Cell Rep. 44, 115558 (2025).

74. Neganova, I., Zhang, X., Atkinson, S. & Lako, M. Expression and functional analysis of G1 to S regulatory components reveals an important role for CDK2 in cell cycle regulation in human embryonic stem cells. Oncogene 28, 20–30 (2009).

75. Liu, L. et al. G1 cyclins link proliferation, pluripotency and differentiation of embryonic stem cells. Nat. Cell Biol. 19, 177–188 (2017).

76. Shariati, S. A. et al. Reversible Disruption of Specific Transcription Factor-DNA Interactions Using CRISPR/Cas9. Mol. Cell 74, 622–633.e4 (2019).

77. Caulier, G., Siblini, J., Sène, L., Mauxion, F. & Séraphin, B. The CCR4-NOT complex: a multifaceted sensor of molecular signals instructing eukaryotic mRNA translation and stability. Nucleic Acids Res. 53, gkaf1401 (2025).

78. Faustova, I. et al. A new linear cyclin docking motif that mediates exclusively S-phase CDK-specific signaling. EMBO J. 40, e105839 (2021).

79. Örd, M., Venta, R., Möll, K., Valk, E. & Loog, M. Cyclin-specific docking mechanisms reveal the complexity of M-CDK function in the cell cycle. Mol. Cell 75, 76–89.e3 (2019).

80. Örd, M. et al. Proline-rich motifs control G2-CDK target phosphorylation and priming an anchoring protein for Polo kinase localization. Cell Rep. 31, 107757 (2020).

81. Shukla, V. et al. HMCES functions in the alternative end-joining pathway of the DNA DSB repair during class switch recombination in B cells. Mol. Cell 77, 1154 (2020).

82. Wu, L. et al. HMCES protects immunoglobulin genes specifically from deletions during somatic hypermutation. Genes Dev. 36, 433–450 (2022).

83. Marton, T. et al. Polymerase theta repairs persistent G1-induced DNA breaks in S-phase during class switch recombination. Nat. Commun. 16, 10536 (2025).

84. Li, S. et al. Microhomology-mediated end joining acts directly on replication forks to repair single-ended double-strand breaks. Mol. Cell 86, 1230–1246.e8 (2026).

85. Baldarelli, R. M. et al. Mouse Genome Informatics: an integrated knowledgebase system for the laboratory mouse. Genetics 227, iyae031 (2024).

86. Zhao, C. et al. A comprehensive human embryo reference tool using single-cell RNA-sequencing data. Nat. Methods 22, 193–206 (2025).

87. Düster, R., Ji, Y., Pan, K.-T., Urlaub, H. & Geyer, M. Functional characterization of the human Cdk10/Cyclin Q complex. Open Biol. 12, 210381 (2022).

88. Schindelin, J., et al. Fiji: an open-source platform for biological-image analysis. Nat. Methods 9, 676–682 (2012).

89. Stirling, D. R. et al. CellProfiler 4: improvements in speed, utility and usability. BMC Bioinformatics 22, 433 (2021).

90. Kõivomägi, M. Purification of cyclin-dependent kinase fusion complexes for in vitro analysis. Methods Mol. Biol. 2329, 95–109 (2021).

91. Zhang, W. et al. Distinct allosteric networks in CDK4 and CDK6 in the cell cycle and in drug resistance. J. Mol. Biol. 437, 169121 (2025).

